# ProteinCLIP: enhancing protein language models with natural language

**DOI:** 10.1101/2024.05.14.594226

**Authors:** Kevin E. Wu, Howard Chang, James Zou

## Abstract

Language models have enabled a new era of biological sequence modeling. However, extracting meaningful sequence-level embeddings from these models remains challenging. In this work, we introduce ProteinCLIP, which applies contrastive learning between a protein’s amino acid sequence and curated text describing its function. ProteinCLIP thus learns to take a pre-trained protein language model’s sequence embedding and refines it produce a function-centric embedding. We show that this embedding space yields sequence representations that enable state-of-the-art performance across a variety of important yet challenging tasks in the study of proteins – from predicting protein protein interactions to accurately detecting homologous proteins despite low sequence similarity. More broadly, ProteinCLIP demonstrates the effectiveness of multi-modal learning in biological contexts, and how such strategies can help isolate key signals from large models and further improve their utility.

## Introduction

Large language models (LLMs) are becoming a powerful technology for studying biological sequences. While LLMs were originally developed targeting natural language applications [1], they have since been successfully adapted to model a wide range of biological sequence modalities spanning protein sequences [2, 3, 4, 5], DNA sequences [6, 7, 8], and RNA sequences [9, 10, 11]. These foundation models for biological sequence have achieved a myriad of state-of-the-art results, including predicting effects of coding variants on protein function [12], effects of non-coding mutations on gene expression [7], protein structure from amino acid sequences [13], protein posttranslational modifications [14], transcriptomic effects of gene knockouts [15], identifying homologous proteins [16], and even generating new proteins [17].

The success of these biological language models is driven, in large part, by their training procedure, whereby random positions in a biological sequence are hidden or “masked” and the language model is tasked with filling in the missing amino acids or nucleotides. When applied across a large corpus of diverse sequences, this relatively simple task learns complex sequence correlations that are informative for much more challenging problems such as protein folding [13] and binding prediction [18, 19]. Despite the success of this approach, it notably ignores the vast knowledge that scientists have gathered over years of studying proteins and their associated functions. Inspired by recent work that has demonstrated that the natural language text describing a biological molecule can be used as an alternative representation to the molecule’s compositional makeup [20], we hypothesize that this scientific knowledge – commonly captured in free text describing protein functions and properties – is complementary to the sequence correlations that have been learned by these biological language models, and that by combining them, we can improve biological language models.

This idea that complementary views of an object can be leveraged to develop richer models has been gaining traction throughout the broader machine learning community. For example, approaches like contrastive learning use pairs of images and their textual descriptions to train models that support rich, flexible query patterns [21], and have more recently become foundational in generative applications as well [22]. Within machine learning works focused on biology, several multi-modal models have been proposed that connect different experimental modalities profiling the same biological system [23, 24, 25]. Although successful, these methods are conceptually narrower, as they bridge very specific measurements that have a direct experimental connection.

In this work, we focus on bridging a specific class of biological language models trained on amino acid sequences, commonly referred to as protein language models (pLMs), with natural language models to create ProteinCLIP. Specifically, ProteinCLIP applies contrastive learning [21] to learn a shared embedding representation between protein amino acid sequences and textual function annotations of those proteins. While classical examples of contrastive learning harmonize embeddings of images and corresponding text describing those images, our contrastive objective harnesses the intuition that the function of a protein is intrinsically driven by its amino acid sequence. In other words, even though amino acids and text may superficially appear dissimilar, they are describing the same system in different vocabularies. In harmonizing amino acids and their emergent function, ProteinCLIP enables downstream applications such as generating sequence-level embeddings that are highly sensitive to deleterious amino acid mutations, state-of-the-art prediction of proteinprotein interactions, and robust homology identification and annotation of unseen sequences. More broadly, ProteinCLIP is a flexible approach that can be applied to harmonize any protein language model with texts describing key aspects of those proteins, and demonstrates that combining knowledge across different modalities is an highly effective, computationally efficient method to focus and distill salient knowledge from pre-trained biological language models.

## Results

### Training ProteinCLIP

ProteinCLIP leverages contrastive learning to learn a shared embedding between amino acid sequences and natural langauge text describing these sequences’ functionality. Architecturally, ProteinCLIP trains two multi-layer perceptron (MLP) “adapter” networks with a single hidden layer that each re-project their respective input protein or text embeddings into a shared embedding space (Figure 1a, S1, see Methods). To learn this shared embedding space, we take after the original CLIP methodology and apply a symmetric cross entropy loss that encourages correct pairings of sequence and text to have similar embeddings while driving other mismatched pairs within the mini-batch to be distant [21]. However, whereas the original CLIP methodology does not leverage pre-trained model weights, ProteinCLIP uses off-the-shelf, pre-trained models for embedding both proteins and natural language without any additional fine-tuning. For proteins, we use the 6, 12, 30, 33, and 36-layer pLMs from the ESM2 family [13] and the ProtT5 pLM [3]. These pLMs produce an embedding corresponding to each amino acid in its input; we average these embeddings to obtain a length-agnostic embedding vector summarizing a protein. For natural language, we use the `text-embedding-3-large` model from OpenAI, whose interface directly returns a length-agnostic embedding for a given text. This approach of training MLP “adapter” networks based on pre-trained embeddings is far more computationally efficient than training entirely from scratch and leverages the significant work that has gone into pre-training foundational models. In addition, since the pre-trained language models themselves are frozen and not updated, our training procedure does not inject additional knowledge into the language model itself, and instead only learns to recombine aspects of what these models have already learned using a relatively small number of parameters (Table S1).

**Figure 1:**
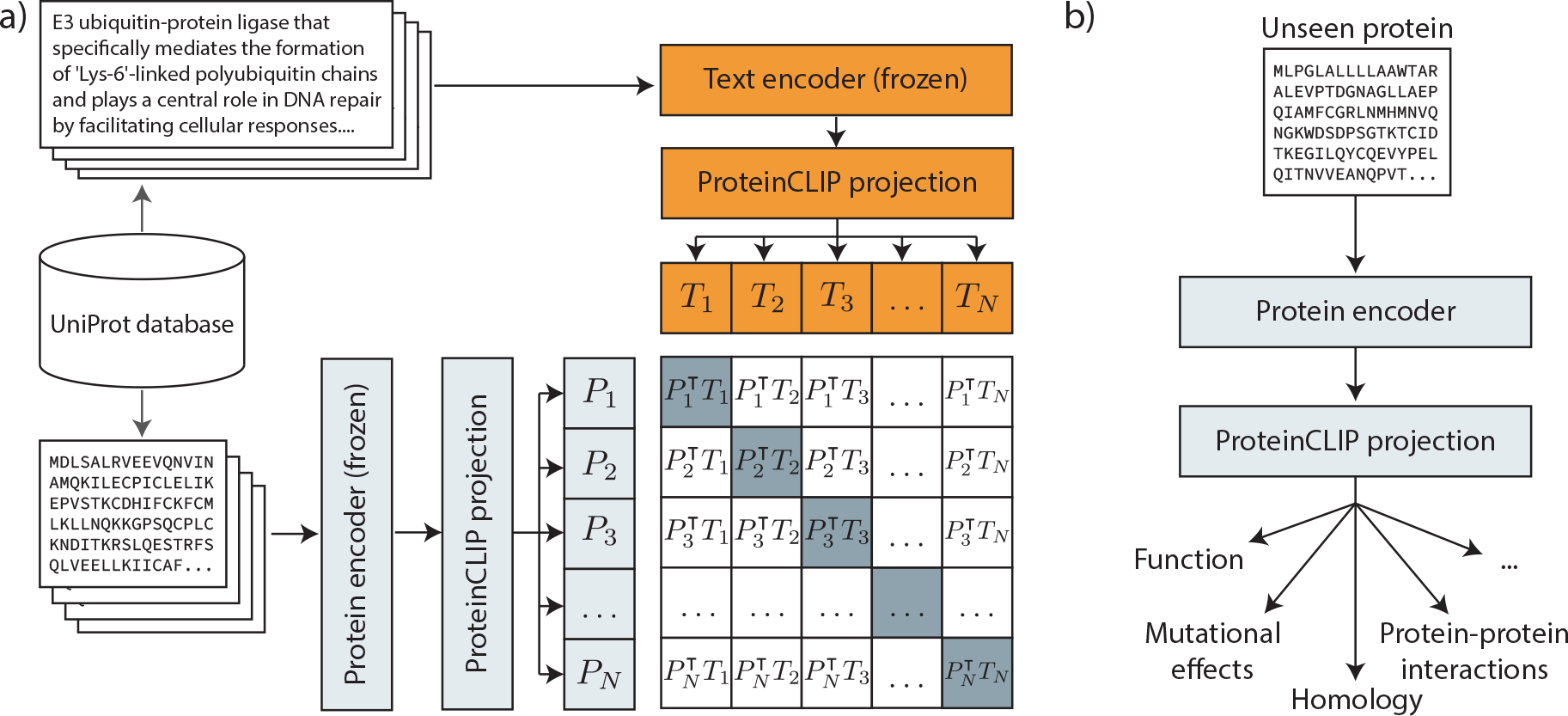
ProteinCLIP performs contrastive learning between a protein’s amino acid representation and a textual description of its function (a). In our work, function texts and amino acid sequences from UniProt are fed through frozen, pre-trained domain-specific language models that serve as encoders. These embeddings are then passed through trainable projection networks to generate the shared embedding space. After training, ProteinCLIP’s protein projection is appended to a pretrained protein language model to embed unseen amino acids in this shared functional space that enables effective prediction of key attributes such as homology and interaction partners (b).

Training data for ProteinCLIP is taken from the UniProt database [26]. We used 465,770 pairs of sequences and function annotation texts (Figure 1) spanning 14,514 different organisms. The functional annotation texts are curated texts describing functional aspects of a protein such as involvement in cellular pathways (Figure S2). These texts do not typically include metadata like the name of the protein in question, the species from which it was observed, or even interaction partners (Table S2). Proteins found across multiple organisms were treated as distinct pairs, as even if they share similar (or even identical) functional descriptions, their corresponding amino acids differed. These function-sequence pairs (Figure S3) were randomly divided into training, validation, and test splits containing 90%, 5%, and 5% of the data (or 419,193, 23,289, and 23,288 examples) respectively. We train 6 distinct ProteinCLIP models – one model corresponding to each of the 6, 12, 30, 33, and 36 layer versions of the ESM2 model [13], and one ProteinCLIP model for the ProtT5 model [3] which contains 24 layers. After training, we used the test set to check that ProteinCLIP indeed projects amino acid embeddings into a space that more closely reflects functional characteristics than raw embeddings (see Supplementary Information, Figure S4). In the remainder of this work, we study how well ProteinCLIP’s embeddings can enable new applications (Figure 1b), particularly in relation to out-of-the-box pre-trained pLMs alone. We examine key tasks spanning major tenets in the study of proteins – the ability to detect cases where small mutations in amino acid sequence meaningfully change global protein characteristics, the ability to identify binding partners for a given protein, and the ability to find homologous proteins that can serve as anchors for better understanding a protein.

### Sensitivity to deleterious missense mutations

To study how well ProteinCLIP has learned to produce robust, generalizable sequence representations closely linked to function, we evaluated how well its embeddings can capture changes induced by protein mutations. Assuming that embeddings reflect function, we intuitively expect that a mutation that causes a large change in protein function should also produce a substantial change in protein embedding. Similarly, mutations with minimal functional impact should produce a minimal change in protein embedding. We evaluated how well various embeddings capture this intuition using a set of mutational scan experimental studies originally curated by Riesselman et al. [27]. This dataset spans 41 experiments evaluating proteins across diverse species and jointly contains 668,339 mutations that each have a corresponding experimental measure of function. For each of these mutations, we embed both the wildtype amino acid sequence and the mutated amino acid sequence and measure the cosine similarity between the two embeddings. For generating embeddings, we evaluate the six aforementioned variants of ProteinCLIP, as well as their underlying pLMs as baselines. To measure performance on each of these 41 datasets, we compute the Spearman’s correlation between measured experimental impact and cosine similarities with respect to wildtype. As lower cosine similarity indicates reduced similarity, and lower function scores indicate greater impact to function, positive correlations indicate that embeddings are more attuned to deleterious mutations. As each experiment profiles functional impact with different measurements, we directly evaluate correlations against experimental values reported in each original study.

We find that ProteinCLIP consistently improves protein embeddings’ sensitivity to deleterious mutations (Figure 2). Evaluating the largest, 36-layer member of the ESM2 model family (Figure 2a), we observe that among the 41 mutational scan experiments, ProteinCLIP’s embedding shifts are more reflective of mutational impact in 37/41 cases compared to using the off-the-shelf ESM2 pLM alone. Furthermore, ESM2 alone counterintuitively produces embeddings that are actually more similar to wildtype for more deleterious mutations in 13/41 mutational screen experiments.

**Figure 2:**
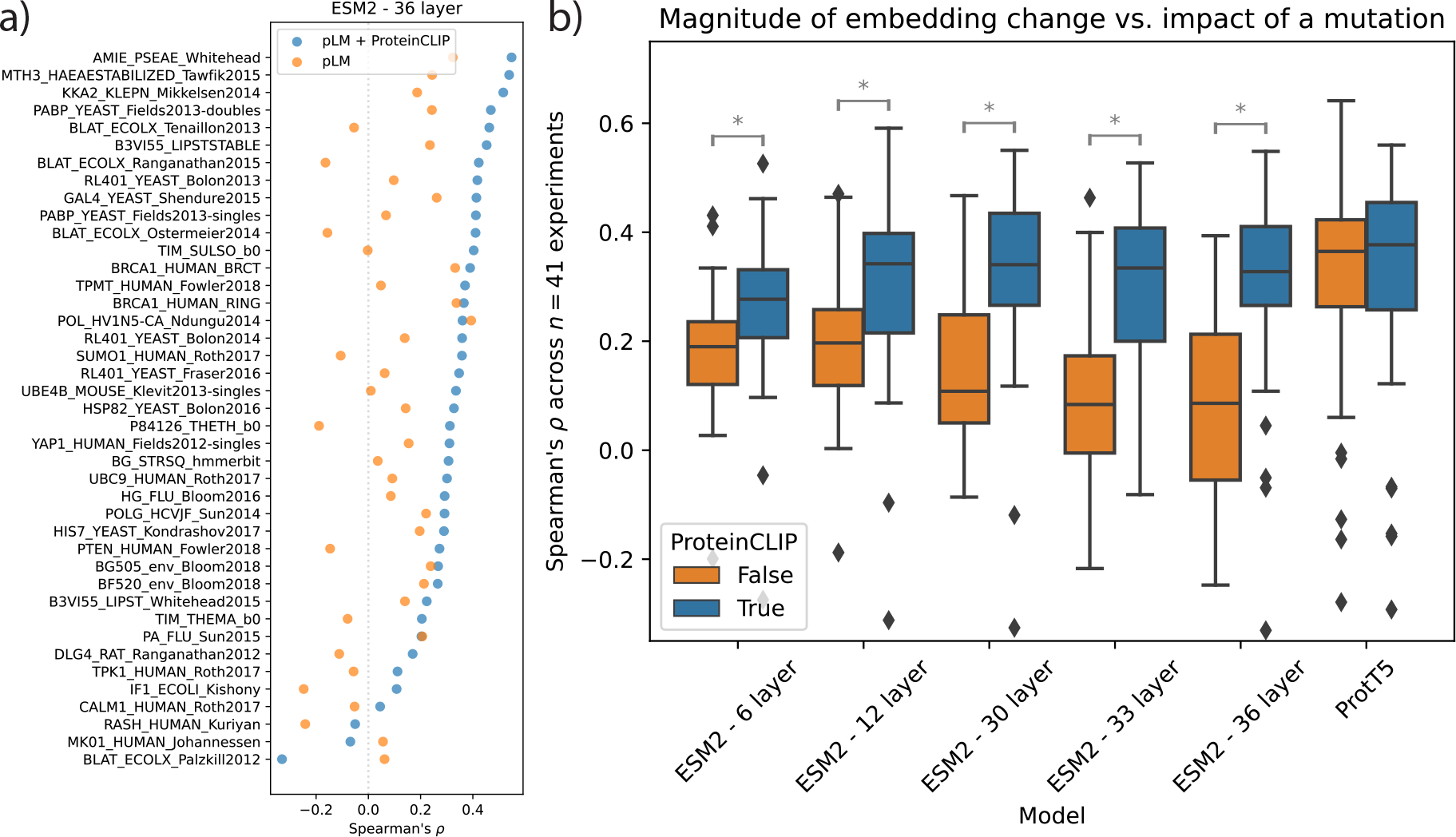
ProteinCLIP and pLM embeddings’ sensitivity to mutations. Each of 41 mutational scan experiments is scored by computing the Spearman’s correlation between experimental function measurements and cosine similarity between wildtype and mutated amino acid sequence embeddings. For the 36-layer ESM2 model (a), ProteinCLIP’s embedding perturbations are better correlated with (x-axis) with functional impact (blue) than the off-the-shelf model (orange) across the experiments (y-axis). Other pLMs show a similar trend (b); boxplots indicate the distribution of Spearman’s correlations for different pLMs (x-axis) with and without ProteinCLIP (blue and orange bars, respectively). Grey asterisk brackets indicate instances where ProteinCLIP’s performance uplift is statistically significant (two-sided Wilcoxon test with Holm-Sidak correction, Figure S5).

In contrast, ProteinCLIP exhibits significantly fewer instances of such erroneous anti-correlated behavior (3/41 cases, *p* = 0.0053, Chi-square test).

These overall trends can be seen when evaluating other pLMs as well. For all models belong to the ESM2 family, ProteinCLIP’s projection yields a significant increase in how well mutational impacts are reflected in embedding shifts (Wilcoxon test with Holm-Sidak correction, Figure 2b), S5). Additionally, we observe that as we scale up the size of the ESM2 models, we do not see consistent improvements in how well their out-of-the-box embedding perturbations mirror functional perturbations. This is contrary to the widely observed trend where larger models tend to learn more effective representations [28]. However, applying ProteinCLIP to these ESM2 models restores the expected gradual increase in these representations sensitivity to deleterious mutations. Indeed, the difference between out-of-the-box ESM2 models and their ProteinCLIP projections appears to grow with larger model sizes. For the ProtT5 model, while ProteinCLIP has higher median Spearman correlation, the difference is not statistically significant. (Figure 2b, S5). This difference may be a consequence of ProtT5’s encoder-decoder architecture [29, 3], which contrasts with ESM’s use of an encoder-only model [13].

Despite these gains, we note that ProteinCLIP itself is not designed to be a strong predictor of mutational effects. There is a rich body of work focused on leveraging pLMs to predict variant effects [30, 2, 12]; these works typically isolate the pLM embeddings or likelihoods at specific mutated positions, which yields a more focused, position-specific signal than averaging across the length of an entire sequence, as we do here. ProteinCLIP is designed to instead provide a sequence-level summary of a protein aggregated across entire sequences with thousands of amino acids. ProteinCLIP’s key contribution is its ability to do so while retaining sensitivity to mutations. This capability is particularly impressive when considering that ProteinCLIP’s training only involves canonical, wildtype sequences; differentiating between deleterious and relatively benign single residue mutations requires generalization at a fine-grained resolution beyond what it is exposed to during training. Furthermore, we find no difference in embedding sensitivity for proteins that were present in ProteinCLIP’s training set compared to those that were not seen during training (Figure S6). Overall, these results suggests that ProteinCLIP’s representations robustly capture functional attributes of proteins; in the following sections, we explore applications enabled by ProteinCLIP’s descriptive sequence-level embeddings.

### Predicting protein-protein interactions

A major goal in the study of proteins is predicting protein-protein interactions (PPIs). Accurate identification of interaction partners for proteins is key both for understanding biological systems [31], and for screening for desirable and undesirable binding affinities when designing novel protein therapies [32]. While several machine learning methods have been proposed to predict whether a given pair of protein sequences interacts or not, recent work has demonstrated that much of the impressive performance typically reported on this task can be attributed to data leakage across training and testing data splits [33]. To combat this, a leakage-free “gold-standard” dataset of human protein-protein interactions has been proposed by Bernett et al. [33] which enforces that there are no overlapping proteins between training, validation, and test sets, and that sequence identity between the sets is capped at 40%. Thus far, methods that are able to achieve non-trivial generalization on this “gold-standard” dataset employ pLMs to generate embeddings for protein sequences [19, 34, 35]. We reasoned that since ProteinCLIP effectively adapts protein embeddings to distill a more function-centric view, it ought to be able to improve PPI prediction.

To predict PPIs, we implemented a simple classifier network that concatenates two sequence-level embeddings (corresponding to two proteins) into a single vector and trains a MLP with a single hidden layer to produce a scalar logit describing predicted binding likelihood (see Methods for additional details). All classifiers we train for this task share this same architecture, isolating the effect of different embedddings. Specifically, we compared two methods for generating these input protein embeddings: as a baseline, embeddings can be directly computed using an off-the-shelf pLM, or they can be additionally passed through our ProteinCLIP method. We perform this baseline-ProteinCLIP comparison using the six different pLM and ProteinCLIP models discussed above (i.e., five ESM2 variants and ProtT5). We observe that for 5 of the 6 pLM evaluated, ProteinCLIP-transformed embeddings yields improvement in PPI prediction performance compared to using the pLM embeddings alone (Figure 3a, S8, Table S3). Moreover, our best-performing model built on ProtT5 followed by ProteinCLIP substantially exceeds the performance of prior methods like Topsy Turvy [34], D-SCRIPT [19], and efforts fine-tuning pLMs for this task [35]. To the best of our knowledge, ProteinCLIP achieves the best performance reported to date on this “gold standard” protein-protein interaction test set.

**Figure 3:**
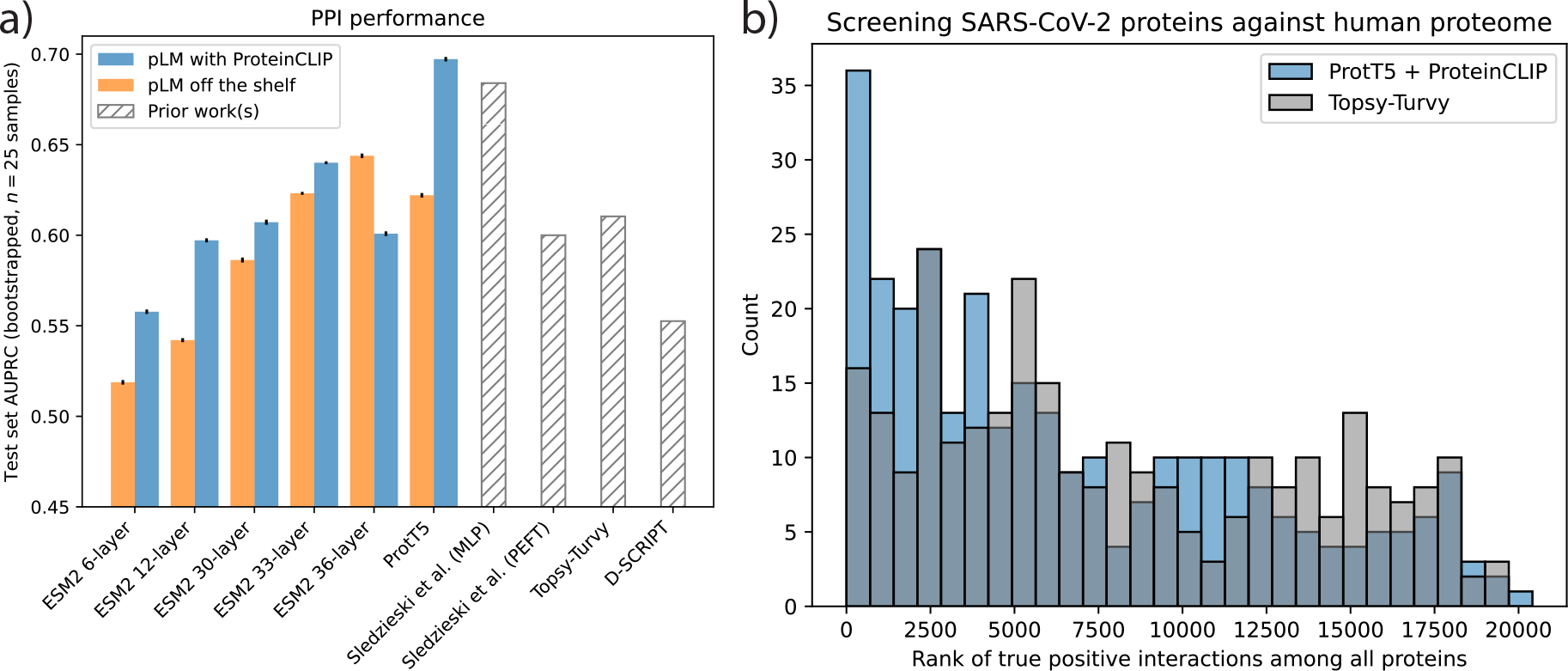
ProteinCLIP and vanilla pLMs’ performance on protein-protein interaction prediction. Panel (a) shows test set area under the precision recall curve (AUPRC, y-axis) when training a MLP to predict PPIs using embeddings generated by an off-the-shelf pLM (orange) or ProteinCLIP (blue). We study the behavior of several models (x-axis), including prior methods (hatched bars). 95% confidence intervals from bootstrapping the test set are shown by vertical black bars. We also evaluated PPI predictors’ ability to prioritize SARS-CoV-2 viral-host protein interactions after training only on human data. We measure the rank of experimentally characterized interactions among a genome-wide screen of all viral proteins against all human proteins for two models: a classifier built on ProtT5 and ProteinCLIP (blue), and a prior method Topsy-Turvy (grey).

While we adhered to the strict data splits when training the PPI classifier network, the ProteinCLIP embedding model itself may have seen some of the test set PPI proteins during training. To account for this, we re-trained ProteinCLIP while enforcing that all proteins in the “gold standard” PPI dataset’s validation and test sets were not used for training ProteinCLIP’s embedding either. We performed this for the 33-layer ESM2 model, as well as for the ProtT5 model, and used the resulting ProteinCLIP projections to train new PPI prediction classifiers (again using the “gold standard” strict data splits). These new leakage-free classifiers have test set performance very similar to our prior results; for example, ProtT5 with ProteinCLIP achieves a test AUPRC of 0.697 originally, and 0.695 after retraining ProteinCLIP with strict holdouts (Supplementary Table S4). This suggests that overlaps between ProteinCLIP’s training and PPI data does not appear to drive large differences in final model performance. This is unsurprising, as the function text we use to train ProteinCLIP does not contain extensive annotations for protein-protein interactions, nor do they contain information that consistently connects protein names and their corresponding sequences (Table S2).

Beyond achieving strong results on this challenging benchmark, we wanted to evaluate the potential of PPI predictors based on ProteinCLIP to be used as a tool for studying novel proteins’ interactions within known systems. As a case study, we examine how well various PPI predictors trained only on human interactions can prioritize experimentally verified viral-host protein interactions between SARS-CoV-2 and human proteins [36]. To evaluate ProteinCLIP in this context, we focus on the ProtT5 model, as it previously exhibited the strongest performance on the “gold standard” human test set. We start by re-training ProteinCLIP itself excluding all SARS-CoV-2 proteins and subsequently re-trained a new classifier on the aforementioned human “gold standard” PPI dataset [33]. This leakage-free ProteinCLIP classifier was then used to score each of the 28 SARS-CoV-2 proteins against every protein in the human proteome (see Methods for additional details). For each viral protein, we find the rank of all experimentally verified human interaction partners [36]. Across the set of 304 total high-confidence viral-host protein interactions, we observe that ProteinCLIP is able to prioritize validated interaction partners despite having never seen SARS-CoV-2 proteins. As a baseline, we also evaluated the best prior PPI predictor with available source code, Topsy-Turvy [34], which we retrained on the same set of human PPIs and performed the same viral-human proteome-wide screen. ProteinCLIP assigns true viral-host interactions significantly higher ranks than Topsy-Turvy (Figure 3b, Wilcoxon test, *p* = 1.39 *×* 10*^−^*^4^). We additionally find that a PPI classifier built directly on ProtT5 embeddings without ProteinCLIP is comparable in performance to Topsy-Turvy (Figure S9), indicating that the improved predictive power can be attributed directly to ProteinCLIP and is not achieved solely with the underlying pLM..

### Homology identification

Identifying proteins from a database that are similar to a query protein is a common task, as it allows scientists to infer key attributes such as function and structure from previously studied proteins. Indeed, identification of such related proteins is also critical for state-of-the-art machine learning protein folding methods like AlphaFold2 [37]. Recently, pLMs have emerged as tool to facilitate computationally efficient retrieval of related proteins, as their embeddings have been shown to contain rich information about a protein’s structure, function, and evolution [16]. Methodologically, this takes the form of embedding the query protein sequence, computing its similarity to a database of proteins and their corresponding (cached) embeddings, and finding the most similar database entries. In the following, we study how ProteinCLIP’s function-oriented protein embedding performs in the context of this task focused on retrieving similar proteins.

For this analysis, we use the pubic CATH dataset (version 4.3.0) of protein domains [38]. As this dataset curates protein domains, there is very little overlap with the set of full-length proteins ProteinCLIP is trained on; only 2% of CATH sequences were present in ProteinCLIP’s UniProt training data (Figure S10). Since low sequence similarity represents the most challenging case for homology identification, we specifically used the S20 CATH dataset, which has a maximum of 20% sequence identity and 60% overlap between any two sequences. This dataset contains 14,942 domains ranging from 9 to 1202 residues in length. These domains are distributed across 5,559 distinct homologous superfamilies, of which 1,543 superfamilies contain more than one domain. For each of the 10,926 domains belonging to these non-singleton superfamilies, we queried the domain against all domains in the CATH S20 dataset, including singletons and excluding itself (i.e., a database of 14,941 sequences). Our query pipeline computes the cosine similarity between the embedding for a query protein against the embeddings for all reference proteins in a database and retrieves the single highest-scoring reference. The embedding itself is produced by either an out-of-the-box pLM, or a pLM followed by ProteinCLIP; we evaluated six different pLMs in this context. We consider a retrieval correct if the retrieved sequence belongs to the same homologous superfamily as the query. This overall setup closely follows that of Schütze et al. [16].

Across all evaluations, ProteinCLIP’s projection uniformly improves performance compared to using a pLM alone (Figure 4a, Supplementary Table S5). Often, the performance improvement is large enough where smaller pLMs coupled with ProteinCLIP are competitive with much larger models. For example, the 12-layer ESM2 model, containing 35 million parameters, coupled with ProteinCLIP (itself containing only 293,408 parameters) achieves a top-1 accuracy of 0.637 – within striking range of the 0.661 accuracy achieved by the much larger 36-layer ESM2 model with 3 billion parameters when used alone. We additionally find that ProteinCLIP’s performance uplift is consistent across queries of different lengths (Figure 4b) – retrieval accuracy with ProteinCLIP is comparable or improved compared to off-the-shelf pLMs at all sequence length intervals. Furthermore, we observe that the performance gap tends to widen with longer inputs. These improvements are also consistent when excluding all sequences with substantial sequence similarity to ProteinCLIP’s training set, ruling out the contribution of data leakage as a driving factor (see Supplementary Information, Table S6).

**Figure 4:**
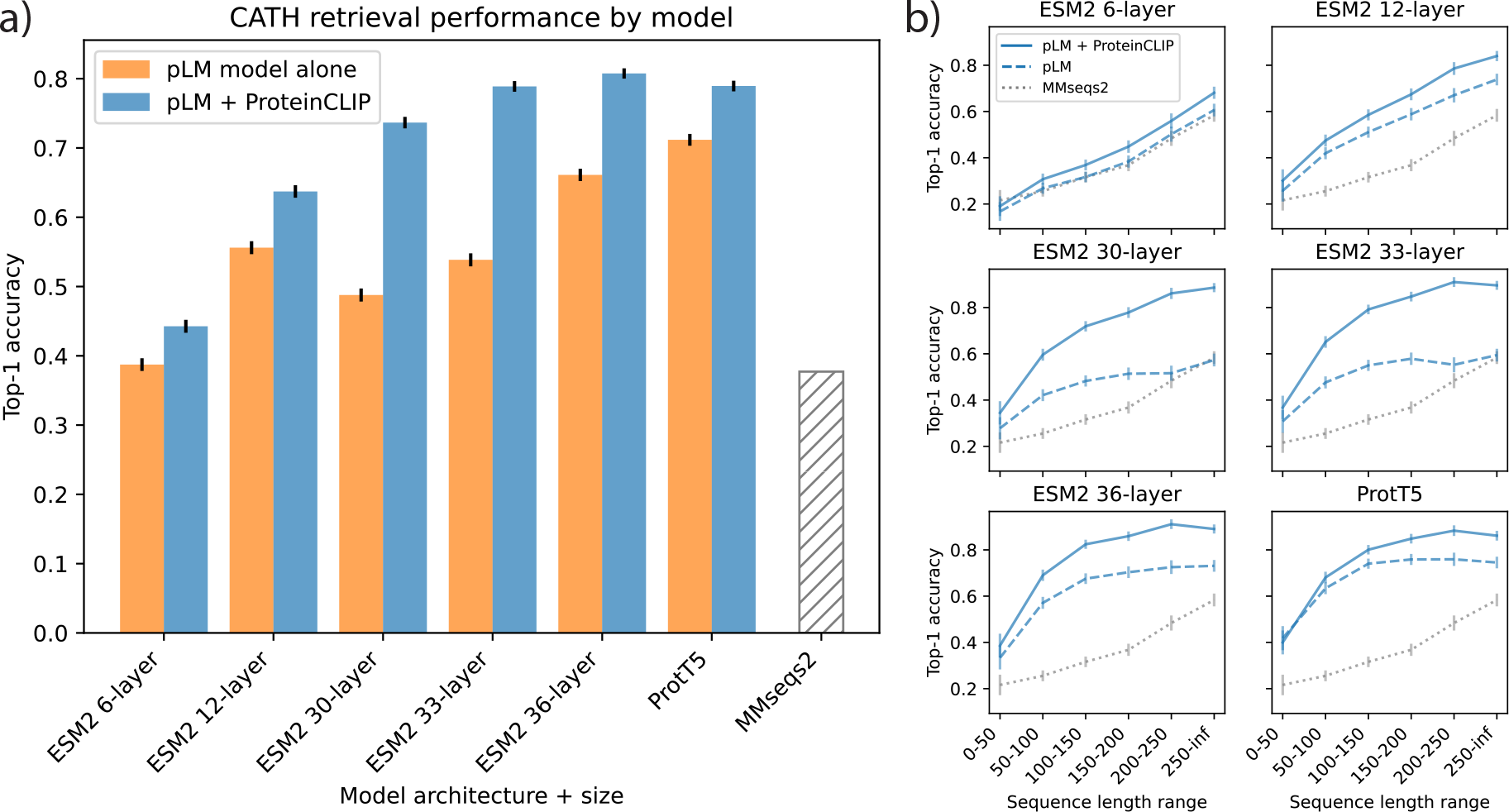
ProteinCLIP compared to pLMs for homology detection among dissimilar sequences. (a) Top-1 accuracy (y-axis) using embeddings from a pLM (orange) or ProteinCLIP (blue) to retrieve a homologous sequence. Different pLMs are shown along the x-axis, black bars indicate 95% confidence intervals. Accuracy of a baseline method MMseqs2 is shown by the hatched bar. (b) Accuracy (y-axis) for query sequences of different lengths (x-axis) for each of the six pLMs evaluated (panels), both with (solid lines) and without (dashed lines) ProteinCLIP. Vertical bars indicate 95% confidence intervals. Grey dotted line indicates performance of MMseqs2.

To better understand how these results compare to conventional methods that use sequence similarity to detect homology, we apply a state-of-the-art retrieval tool, MMseqs2 [39], to this same CATH S20 dataset (see Methods for additional details). MMseqs2 performs similarly to the smallest pLM we evaluated, the 6-layer ESM2 model (Figure 4a, Supplementary Table S5), with an accuracy of 0.377 compared to 0.387 for the latter. This performance gap is consistent across sequences of different lengths (Figure 4b) and grows as we use larger pLMs and when we apply ProteinCLIP. These results suggest that pLMs and especially ProteinCLIP can greatly improve our ability to detect homologous proteins with low sequence similarity.

## Discussion

ProteinCLIP is a contrastive approach that harmonizes the amino acid embeddings produced by a protein language model, and textual embeddings of corresponding protein function descriptions generated by a natural language model. ProteinCLIP does not perform any fine-tuning of the language models it is built off of, and thus cannot “inject” additional information into these models. It instead learns to distill a more succinct, effective representation of function by reconfiguring the embedding that a protein language model has already learned. Despite its architectural simplicity, ProteinCLIP is highly effective at distilling amino acid embeddings towards a more function-centric view. We demonstrate that ProteinCLIP is able to generalize to unseen amino acid sequences and achieve state-of-the-art results in predicting those proteins’ global response to mutations, identifying protein interaction partners, and finding homologous proteins with low sequence similarity. We surmise that these tasks in particular benefit from aligning sequence and function embeddings, as they require a global understanding of a protein and its function. For example, after isolating aspects of pLM embeddings informative for overall function, examining changes in these function-oriented embeddings is more likely to reveal changes in function than the original raw embeddings. Similarly, homologous proteins are likely to perform similar functions, and as such a function-focused embedding is likely to amplify signals relevant for homology. Beyond strong predictive performance, these improvements to protein annotation pipelines can serve as a basis for quickly studying and understanding biological systems *in silico* (as we demonstrate using SARS-CoV-2 proteins), and even for screening novel protein designs for desirable behaviors at scale.

Despite significant gains across a variety of tasks, ProteinCLIP has limitations as well. Since ProteinCLIP is fundamentally tuned towards protein function, it does not enable performance gains in tasks that are not directly related to protein function. For example, we evaluated ProteinCLIP’s capacity to improve prediction of interactions between small molecules and proteins using a method closely inspired by Singh et al. [40], but found no performance gains, likely due to the fact that such interactions are more fundamentally driven by structural knowledge rather than functional annotation. This limitation suggests a line of future work, whereby we incorporate additional modalities or additional text annotations in the contrastive learning objective to learn a more general embedding representation. Another limitation of our ProteinCLIP methodology is its reliance on average pooling an embedding before applying contrastive learning. Prior works have pointed out limitations of average pooling [41]. While we explored alternative methods such as training an attention-based aggregation mechanism to pool along a sequence in a more nuanced, context-aware fashion, we found that doing so resulted in an overall network that was too large to train effectively given our hardware constraints. Future work that builds contrastive approaches similar to our work, but with less naive pooling strategies may yield even greater performance gains.

ProteinCLIP is a useful resource for the scientific community, both by advancing performance across a number of key protein annotation tasks, and by representing a broader view that multimodal modeling in biological contexts is an effective strategy. Indeed, while self-supervised pretraining strategies have proved incredibly successful for modeling biological and natural text sequences, ProteinCLIP demonstrates that judiciously combining these strategies in a multi-modal fashion allows us to extract even more from these already powerful models. More importantly, ProteinCLIP demonstrates that doing so does not necessarily require large computational investments that training ever more larger language models entails, and that we are far from reaching the full potential that these models offer.

## Methods

### Language models

All protein language models used in this work are open-source and freely available. We specifically use two families of pLMs – the evolutionay scale modeling (ESM) family [13], and the ProtT5 family [3]. All pLMs used in this work were pre-trained on the same UniRef50 dataset [42] (though exact data splits vary), so differences between models are primarily architectural. For ESM models, we use the following pre-trained checkpoints available through their codebase at https://github.com/facebookresearch/esm:

- 36-layer model: `esm2_t36_3B_UR50D`
- 33-layer model: esm2_t33_650M_UR50D
- 30-layer model: esm2_t30_150M_UR50D
- 12-layer model: esm2_t12_35M_UR50D
- 6-layer model: `esm2_t6_8M_UR50D`

For ProtT5, we use the encoder-only, half-precision model as it was indicated by the original authors to produce better embeddings; the model is available at the HuggingFace hub at https://huggingface.co/Rostlab/prot_t5_xl_half_uniref50-enc.

All protein models were frozen (i.e., not fine-tuned or trained). We also do not re-train pLM models to remove data leakage as we do for the ProteinCLIP projection itself, as re-training these pLMs would be prohibitively time-consuming and resource intensive. Sequence level embeddings were obtained by taking an amino acid sequence and passing it through each model, which yields an embedding for each position in the input sequence. We average all embeddings along the sequence length dimension (excluding special characters indicating beginning/end of sequence) to get a fixed-size embedding for each protein. We only showed these models full proteins in their entirety and did not perform chunking or splitting as some other works have explored [12]; proteins too long to fit in memory were discarded from training and downstream analysis (see below). We explored the possibility of fine-tuning these models in order to train the initial “beginning of sequence” special tokens to produce a sequence summary embedding, but the size of these models did not fit into GPU memory on our available hardware.

For natural language modeling, we use OpenAI’s public facing API to access their embedding model `text-embedding-3-large`. This API yields a vector summarizing the entirety of the text given to it.

### ProteinCLIP architecture and training

ProteinCLIP consists of two projection networks that each take an input embedding and re-projects it into a shared embedding space. These projection networks are simple multi-layer perceptron (MLP) networks with a single hidden layer that are constructed as follows. A MLP that processes an input of dimensionality *d* first projects into a *d*-dimensional hidden space followed by GELU activation [43] and a LayerNorm [44]. Secondly, this *d*-dimensional hidden space is projected into a *d*_shared_-dimensional shared embedding space; all outputs in this shared embedding space are normalized to unit length before being returned by the network. For all ProteinCLIP models shown in this work, we set *d*_shared_ = 128; this represents a significantly smaller embedding size than all language models used in this study (Table S1).

This specific architecture – particularly the number of hidden layers – was selected based on performance on the validation set. We used the training procedure described below and evaluated models with no hidden layers and models with 1, 2, or 3 hidden layers (Figure S1). We observe that models with 2 or 3 hidden layers appear to overfit to training set, as their validation loss plateaus and even increases towards the end of training. The model with a single hidden layer performs similarly to the model with no hidden layers (i.e., linear transformation). We elect to use the single hidden layer model to slightly increase expressiveness without any apparent loss in validation set performance.

ProteinCLIP is trained using the contrastive loss function introduced by Radford et al. [21]. Specifically, given a batch of *N* examples, we compute all pairwise cosine similarities between the outputs of the two MLP networks, yielding a *N × N* matrix of logits. This logit matrix is then scaled by a learnable temperature parameter *t* by multiplying with *e^t^*. We then calculate a standard cross entropy loss whereby all *N* entries on the diagonal are considered positive examples, and all *N* ^2^ *− N* off-diagonal examples are considered negative examples. We do this along both axes of the matrix and average the two losses as our final loss.

ProteinCLIP is trained with the AdamW optimizer [45] and a learning rate of 1 *×* 10*^−^*^4^ over 300 epochs. When training ProteinCLIP itself, we do not use the validation set to choose a best-performing checkpoint, nor do we perform early stopping based on validation set performance – the validation set is primarily used to develop and tune the architecture (see above). Given that contrastive learning tends to benefit from larger batch sizes [46, 21], we use a batch size of 20,480 to train all ProteinCLIP models, except for models built on top of the 36-layer ESM2 model. These models use a slightly smaller batch size of 16,864 to fit in GPU memory as a consequence of its larger embedding dimension.

It is also important to note that these ProteinCLIP projection MLP networks are specific to each base protein language model. Thus, when we compare ProteinCLIP models across different pLMs, we are not comparing the same exact adapter MLP with the same weights, but are comparing MLPs that are designed and trained in the same way. Notably, we also do not perform any tuning of ProteinCLIP or its training procedure to tailor it to different base protein language models. The original design of ProteinCLIP was prototyped using the 33-layer ESM2 model, and the same architectural choices and training procedure were applied for other protein language models, with only *d* varying between models as a point of necessity. It is possible that additional improvement can be gained by tuning the architecture towards specific models.

### Contrastive dataset

Training data for ProteinCLIP is taken from the UniProt database [26]. We downloaded all UniProt records as of February 15, 2024. Records were downloaded in the form of fasta files for sequences, and `.dat` text files for function records. Records with no functional annotation texts were discarded. Records whose amino acid sequences exceeded 5800 amino acid residues in length were discarded due to GPU memory constraints. Although different pLMs have different memory requirements, this sequence length was chosen based on the largest model (ESM2, 36 layer) and enforced for all models such that all models see the exact same examples. 570,037 records remained spanning 14,514 different organisms (Table S2). Data splits into train, validation, and test splits are created randomly, and are identical across all training runs across different models except when explicitly indicated otherwise (e.g., when holding out PPI training data from ProteinCLIP training, holding out SARS-CoV-2 records, etc.).

### Mutational screens

Mutational screen data were taken from many experimental works [47, 48, 49, 50, 51, 52, 53, 54, 55, 56, 57, 58, 59, 60, 61, 62, 63, 64, 65, 66, 67, 68, 69, 70, 71, 72, 73, 74] that were originally aggregated by Riesselman et al. [27]. These mutational screens introduce one or more mutations to a protein and measure the resulting impact to protein function, yielding labeled pairs of mutations and a continuous value profiling their functional impact. As different mutational screen experiments often measure function with different units and scales, functional impact values are not directly comparable between datasets. As such, we focus on studying variance within each dataset rather than between datasets.

Mutational screens were evaluated based on the intuition that an ideal embedding function should be reflective of mutational impact – i.e., mutations measured to be highly disruptive to protein function should cause large changes in embeddings, whereas mutations with minimal functional impact should cause relatively small changes in protein embedding. We use cosine similarity to measure embedding similarity. Within each dataset, we take each mutated sequence, embed it using either an off-the-shelf pLM, and additionally using ProteinCLIP, and measure the cosine similarity between the mutated sequence and the wildtype sequence. We take these values and compute the Spearman’s correlation between cosine similarity and experimentally measured impact. We choose Spearman’s correlation in this context because functional scores may not always be linear.

We evaluate the difference in performance between embedding methods by using Wilcoxon’s signed-rank test to compare the Spearman’s correlation produced by different methods using the above approach. In cases where we perform a statistical test for each pair of base pLM and its ProteinCLIP extension, we perform multiple hypothesis correction using the Holm-Sidak method.

### Protein-protein interaction classifier and dataset

Our protein-protein interaction model is a simple MLP with a single hidden layer. The network takes as input the concatenation of two protein embeddings, and projects them to a 128dimensional hidden layer, followed by GELU activation and layer normalization. This hidden layer is then followed by a linear projection to a 1-dimensional scalar logit corresponding to the network’s binding prediction. This network is trained for 200 epochs with a batch size of 4096 using the AdamW optimizer, and a learning rate of 2 *×* 10*^−^*^4^. As 200 epochs provides sufficient gradient steps to overfit to the training data (Supplementary Figure S7), we used the validation set in the “gold standard” dataset to choose a checkpoint that yielded the highest validation set AUPRC, and used the corresponding model weights to evaluate on test set.

During both training and evaluation, we excluded protein pairs that involved one or more proteins that exceeded our specified length cap of 5800 sequences, or proteins that no longer exist in the official UniProt set. For the test set, there are six such proteins – UniProt identifiers Q09666 (length of 5,890), Q6RUI8 (deprecated), Q7Z4U5 (deprecated), Q8TAB7 (deprecated), Q8WV35 (deprecated), Q8WZ42 (length of 34,350). The exclusion of these identifiers has a minimal impact on test set composition, reducing the size of the test set by a mere 13 examples from 52,048 to 52,035 examples.

We explored several alternative methods of combining the input embeddings aside from concatenation, such as to be more robust to the order in which a pair of proteins is presented. We evaluated the order-agnostic approach of summing together the two input embeddings as in Chen and Zou [20], as well as other methods like taking the difference between the embedding vectors, but saw no meaningful change in validation set performance so did not pursue these methods further. We additionally evaluated a data augmentation approach whereby the order of protein pairs are shuffled when training the PPI classifier; we did not observe meaningful changes in validation set performance with this approach either. We elected to use the concatenation of embeddings, as it was the simplest, most direct approach.

Confidence intervals are obtained using bootstrap analyses. We resampled the “gold standard” test set (with the 13 aforementioned records excluded) 25 separate times and evaluated our trained models on the resampled test set to <u>obtai</u>n 25 estimates of test set performance. Confidence intervals are then determined by 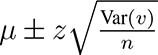 where *µ* denotes the mean across *n* test set performance metric values *v*, and *z* = 1.96. Var denotes a standard variance calculation.

Performance metrics for fine-tuned and low-rank adaptation models from Sledzieski et al. [35] were taken directly from the original manuscript; we did not evaluate this method ourselves as there was no publicly available code at the time of this writing. We re-trained and evaluated Topsy-Turvy [34] and D-SCRIPT [19] using the original authors’ code (https://github.com/samsledje/D-SCRIPT) based on instructions provided by Bernett et al. [33] at https://github.com/biomedbigdata/data-leakage-ppi-prediction. Due to GPU memory constraints, we limited sequences used for Topsy-Turvy and D-SCRIPT to a maximum of 1500 amino acids during training and evaluation, which aligns with the authors’ recommendation for maximum sequence length. This is a longer cap than used by Bernett et al. [33], so final performance numbers differ slightly when comparing benchmark results between these two works.

When measuring performance on SARS-CoV-2/human protein interactions identified by [36], we use the following procedure. For each of 28 SARS-CoV-2 viral proteins (sequences provided by Gordon et al. [36]), we predict an interaction score for all human proteins that fit in GPU memory. For ProtT5 and ProteinCLIP models with a limit of 5800 amino acids, 20414/20428 human proteins are retained, and 19484/20428 for D-SCRIPT and Topsy Turvy which share a length limit of 1500. For all models, proteins are given in the order (viral protein, human protein). Ranks are calculated by taking all the human proteome predictions corresponding to a given SARSCoV-2 protein, and calculating the rank of true human protein interaction partners. This process is repeated for each SARS-CoV-2 proteins and its known human interaction partners. Statistical testing comparing different models excludes any human proteins that are not present in one of the models’ outputs due to aforementioned length restrictions.

### Homology detection

We frame homology detection as a retrieval task and follow a standard retrieval pipeline. We embed each query amino acid sequence using either a protein language model, or a protein language model followed by its corresponding ProteinCLIP model. These embeddings are then used to search a database of proteins (whose embeddings are generated using the same method) using cosine similarity to measure the similarity between the query and each database protein. Cosine similarity is a standard metric for embedding comparison, and is defined between two vectors *u, v* as 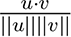; values are bound between [1, -1] with higher values indicating greater similarity. In all cases, we retrieve the single best match with the highest cosine similarity.

Confidence intervals for homology detection results are com<u>puted</u> using the expression for confidence intervals in a binomial distribution, namely 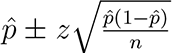 where *p*^ denotes the observed accuracy of the classifier, *z* = 1.96, and *n* indicates the number of samples. We use this to compute confidence intervals for top-1 accuracy as a whole, and for top-1 accuracy broken down by sequence length bins.

We compare our embedding-based retrieval homology detection methodology against a state of the art homology detection method based on sequence comparisons, MMseqs2 [39]. We use a pre-compiled binary, version hash `d4841a8efad066e9758b6626cc64c5ef5ee53055`. Searching was done by taking the CATH dataset sequences, creating a database of all 14,942 domains, and querying the database against itself with an e-value cutoff of `-e 10000`, sensitivity of `-s 7.5` and a maximum sequence limit of `-max-seqs 300`; all other parameters were left at their default values. Results were parsed to obtain the best match for each domain belonging to a non-singleton superfamily that was not itself, as determined by the highest bit score.

### Software packages

ProteinCLIP and all additional networks presented in this work are trained using PyTorch [75] and PyTorch Lightning [76]. Additional analyses were performed using numpy [77], scipy [78], statsmodels [79], and pandas [80]. Plots were made using matplotlib [81] and seaborn [82].

### Model and data availability

As a resource for the community, we make the ProteinCLIP code available on GitHub at https://github.com/wukevin/proteinclip. In addition, trained ProteinCLIP “adapter” networks are available for all the protein language models we examine in this work, namely the 6, 12, 30, 33, and 36-layer ESM2 models, as well as the ProtT5 model; plase refer to our code for instructions for loading and using these models.

For reproducibility, we make all data used to train ProteinCLIP publicly available at the following Zenodo link: https://zenodo.org/doi/10.5281/zenodo.11176862. This includes embeddings of amino acids for the models we studied in this work, as well as text embeddings for their corresponding function descriptions. We also deposited files containing the source amino acids and raw text for the aforementioned embeddings. All datasets used for analysis of ProteinCLIP are publicly available and can be found via their respective original works; when possible, our public code release also provides code for easily loading and working with these datasets.

## Acknowledgements

Author contributions

## Supplementary Information

### Test set validation

After training ProteinCLIP, we wanted to understand how well it performed at the task it was explicitly trained to do: bridging amino acid sequences and their associated functions. We reasoned that one measure of this would be through correlations in pairwise embedding space distances. In particular, if two proteins’ functions are similar as measured by closeness in text embedding space, we expect their protein amino acid embeddings to ideally be close – and vice versa. As a baseline, we compared all pairwise cosine distances between function text embeddings and amino acid embeddings derived from the 33-layer version of ESM2 for all records in the test set, and found a positive Pearson’s correlation of *r* = 0.216. We then took the amino acid embeddings and fed them through ProteinCLIP’s projection network, performed the same cosine distance calculation and correlation analyses using these re-projected protein embeddings, and found an increased Pearson’s correlation of *r* = 0.250 (Figure S4a, b). This increase suggests that on the test set, ProteinCLIP’s contrastive objective learns an embedding that is more reflective of function than the protein language model itself learns.

We reasoned that this relatively small global improvement in pairwise correlations would be larger if we focused on challenging pairs that are functionally similar but dissimilar by sequence. We explored this by focusing on pairs of records in the test set with both low sequence similarity and high function similarity. Specifically, these pairs of test set sequences have pairwise protein embedding distances in the highest 2nd percentile, and function embedding distances in the lowest 2nd percentile. For this set, we find that the out-of-the-box protein embeddings’ pairwise cosine distances have no meaningful correlation with functional similarity (Pearson’s *r* = *−*0.058) whereas ProteinCLIP’s projected version of these embeddings achieves a Pearson’s *r* = 0.270 (Figure S4c, d). We observe similar results for the top 5% intersection (i.e., intersecting bottom 5% of pairwise protein similarities, top 5% of function similarities, Figure S4e, f). This suggests that the aforementioned small global improvement in pairwise distance correlations between function text and amino acid embeddings is primarily driven by sizable improvements for these difficult cases. These results indicate that on the unseen test set, ProteinCLIP models learn an embedding that is more reflective of functional relationships than raw pLMs alone.

### Homology identification with sequence similarity cutoffs

While the CATH S20 dataset has minimal overlap with the UniPort sequences used to train ProteinCLIP, there is still a small percentage of sequences that either overlap directly or bear significant sequence simialrity (Figure S10). We wanted to ensure that the strong results presented in the main body of this work were not an artifact of train/test set bleed, and as such performed a homology retrieval evaluation while holding out these similar and overlapped sequences. Specifically, we took the same CATH S20 dataset, and removed all instances where a sequence was present in the UniProt set, or had a 20% or lower normalized edit distance to the closest UniProt sequence. Normalized edit distance is defined as the Levenshtein edit distance divided by the length of the CATH sequence in question. This resulted in a dataset of *n* = 13, 227 total CATH sequences spanning 5176 homologous superfamilies, of which 1371 superfamilies are non-singletons with at least two sequence members. As in the main text, we take the 9422 sequences belonging to these non-singleton families and identify the top, non-self hit along all other sequences (including singleton families).

We observe a slight but consistent decrease in performance across both pLMs and pLMs combined with ProteinCLIP in this reduced sequence similarity setting. For pLMs, we see drops in accuracy from 0.002 to 0.012, whereas for ProteinCLIP, we see drops in accuracy ranging from 0.012 to 0.020. In all cases, both modeling approaches have slightly poorer accuracy, and ProteinCLIP has a small but consistently larger reduction in accuracy. Importantly, the trend that ProteinCLIP vastly outperforms its off-the-shelf pLM counterpart is still true in this setting, indicating that train/test bleed does not drive over-inflated performance values on this task.

## Supplementary Figures and Tables

**Supplementary Figure S1:**
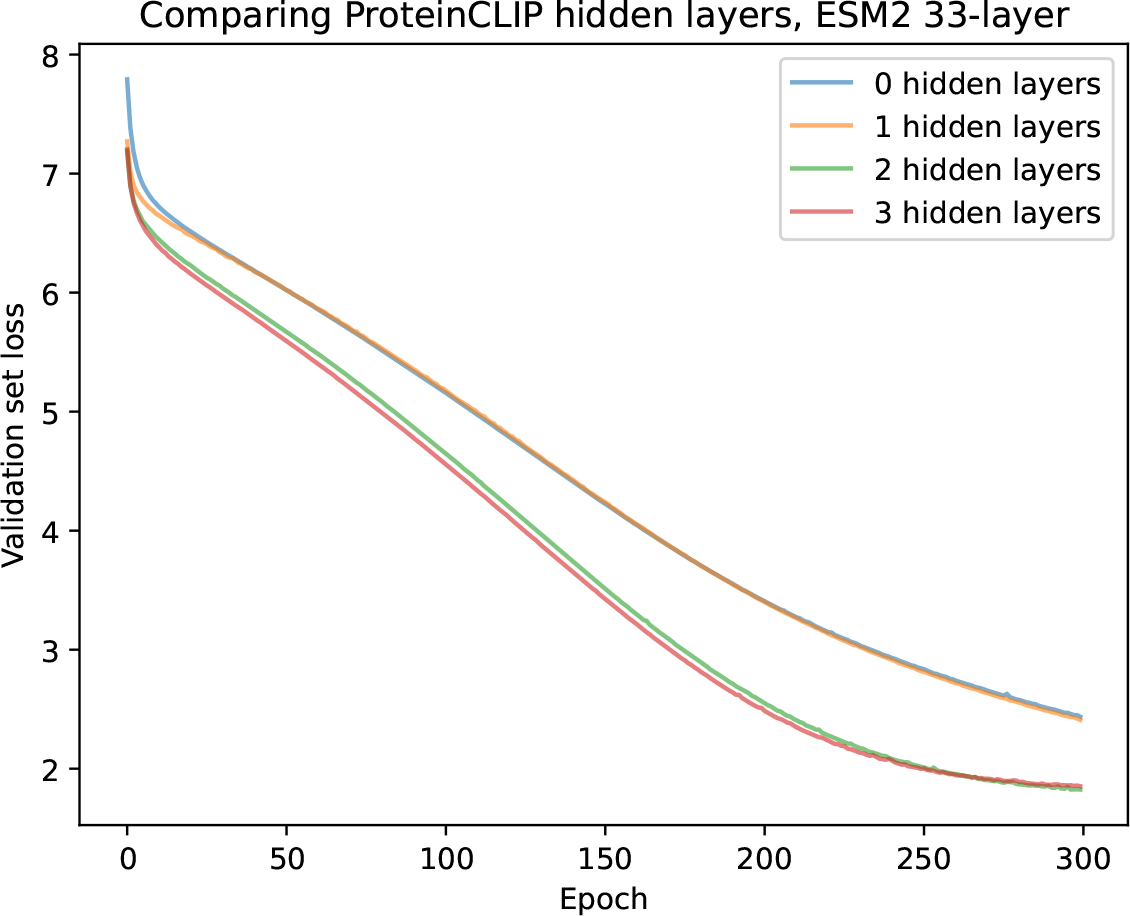
Validation loss curves when training variants of ProteinCLIP with differing numbers of hidden layers (colors). 0 hidden layers indicates a linear projection from input to output embedding. All models are trained with a consistent batch size of 8192; all other training configurations are the same as are described in methods.

**Supplementary Figure S2:**
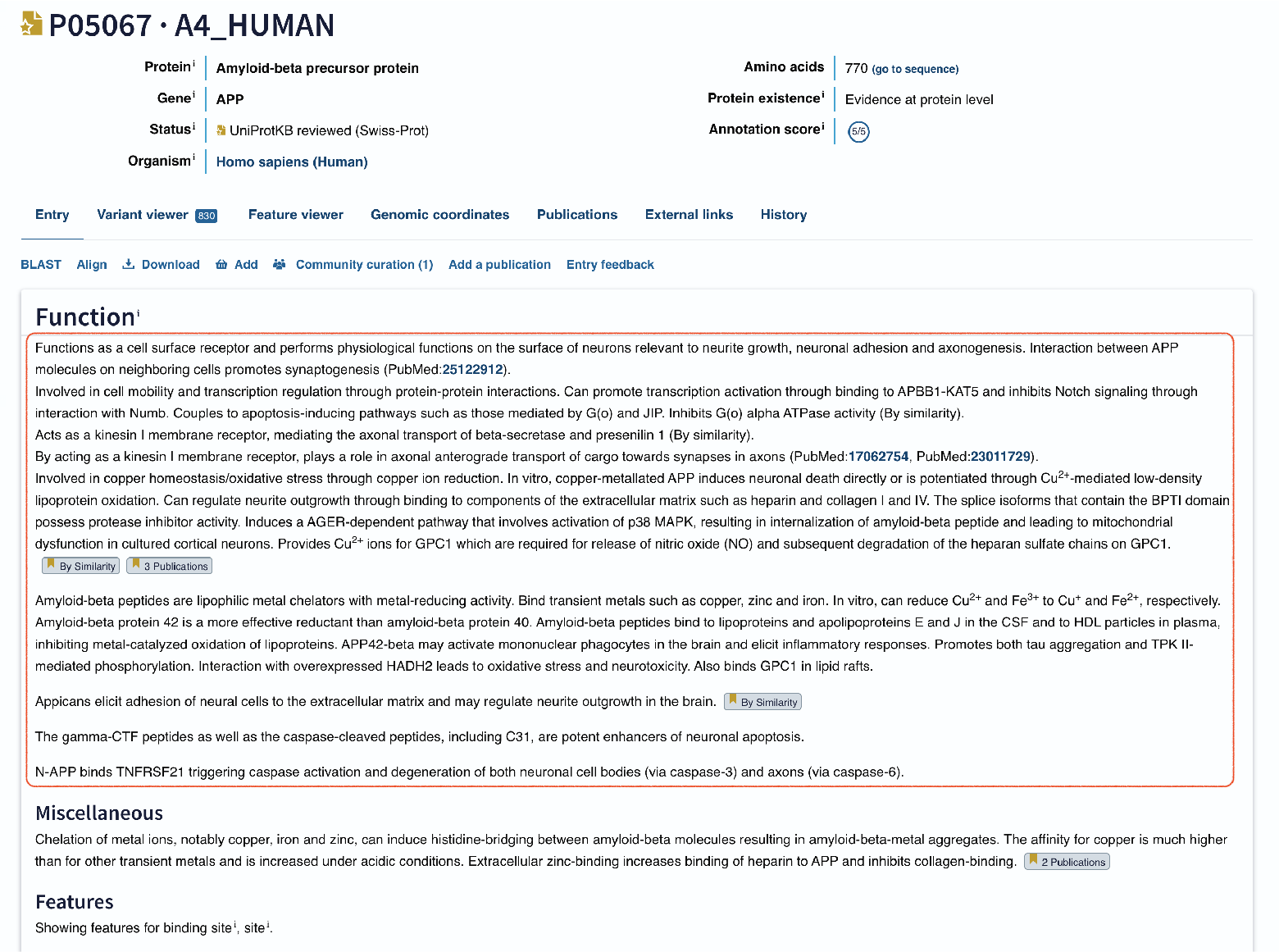
Screenshot of example record P05067 with function text used to train ProteinCLIP highlighted in red box. Function text is parsed from a `.dat` text file rather than parsing the web view (see Methods). References are removed prior to generating text embeddings.

**Supplementary Figure S3:**
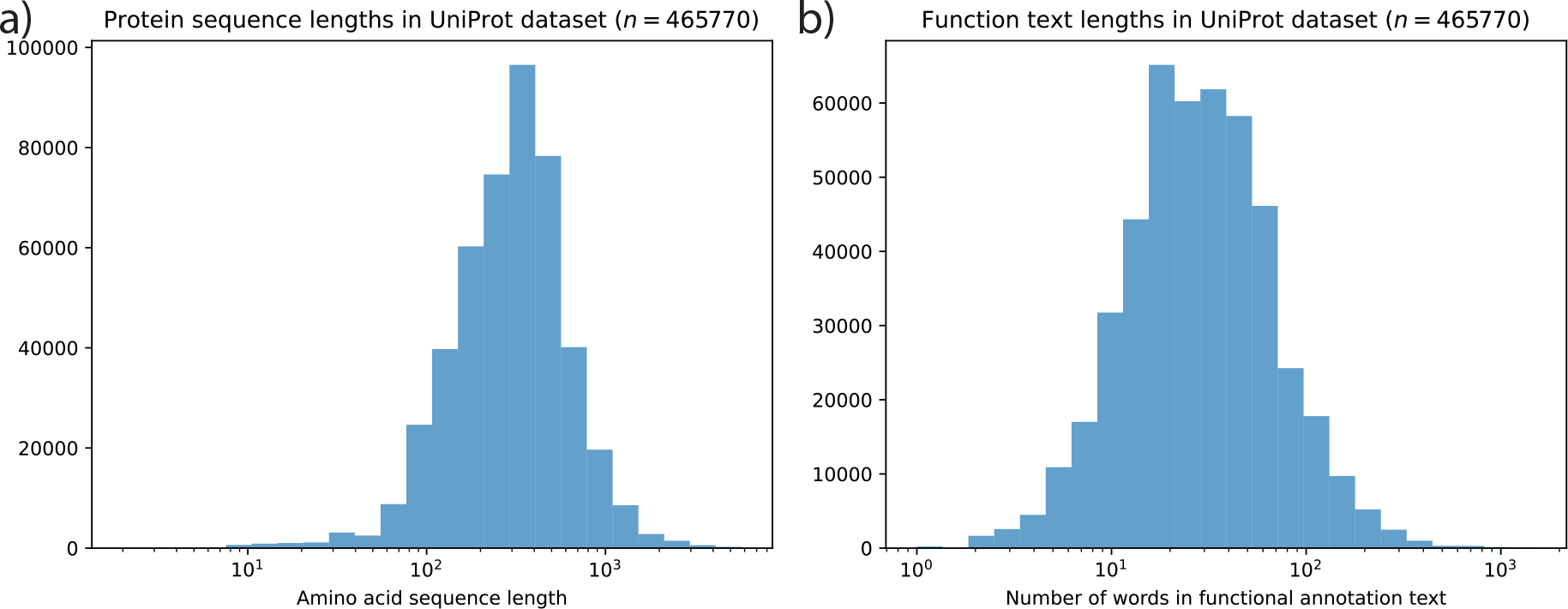
Lengths of amino acid sequences and function texts in UniProt data used to train ProteinCLIP. Panel (a) shows the distribution of amino acid lengths (x-axis, log scaled), which are capped at 5800 residues due to hardware memory restrictions. Amino acid lengths range from 2-5762 residues, with a median length of 307 residues. Panel (b) shows the distribution of function description lengths in number of words (x-axis, log-scaled). Functions range in length from 1-1508 words, with a median length of 28 words.

**Supplementary Figure S4:**
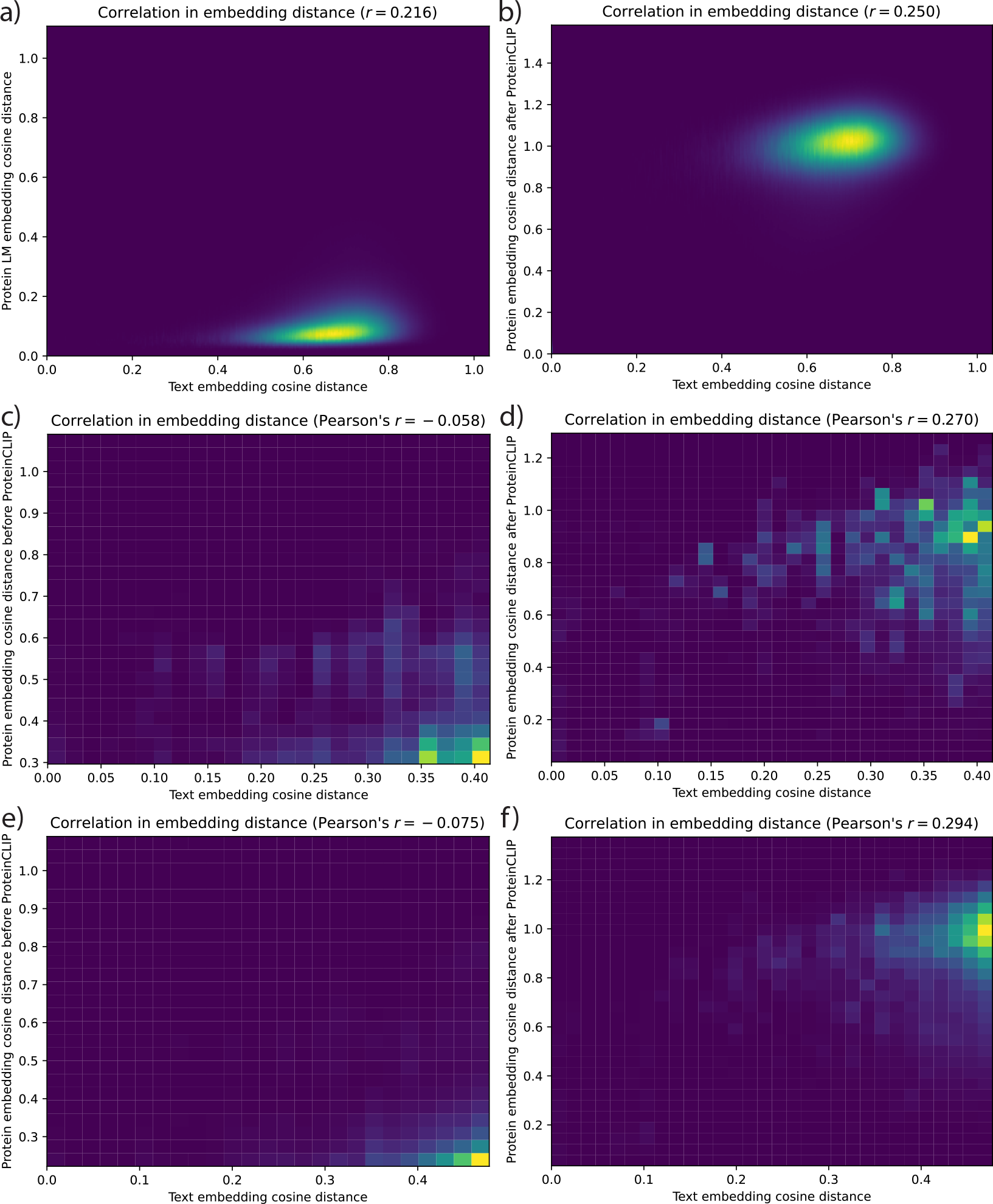
Pearson’s correlation between pairwise cosine distances of two proteins given function text embeddings (x-axis) and amino acid embeddings from the 33-layer ESM2 model (y-axis). An amino acid embedding that perfectly recapitulates functional text produces a perfect correlation of *r* = 1.0. Panels (a, c, e) show how well the pLM performs in its pre-traiend form without any additional modification. Panels (b, d, f) show performance with ProteinCLIP applied. Panels (a, b) show performance when comparing pairwise cosine distances across the entire test set of UniProt entries, panels (c, d) and (e, f) focus on more challenging pairs that have low sequence similarity and high functional similarity, respectively. These challenging pairs are selected by intersecting the top 2% of functional similarity and bottom 2% of amino acid embedding similarity, and top 5%, respectively.

**Supplementary Figure S5:**
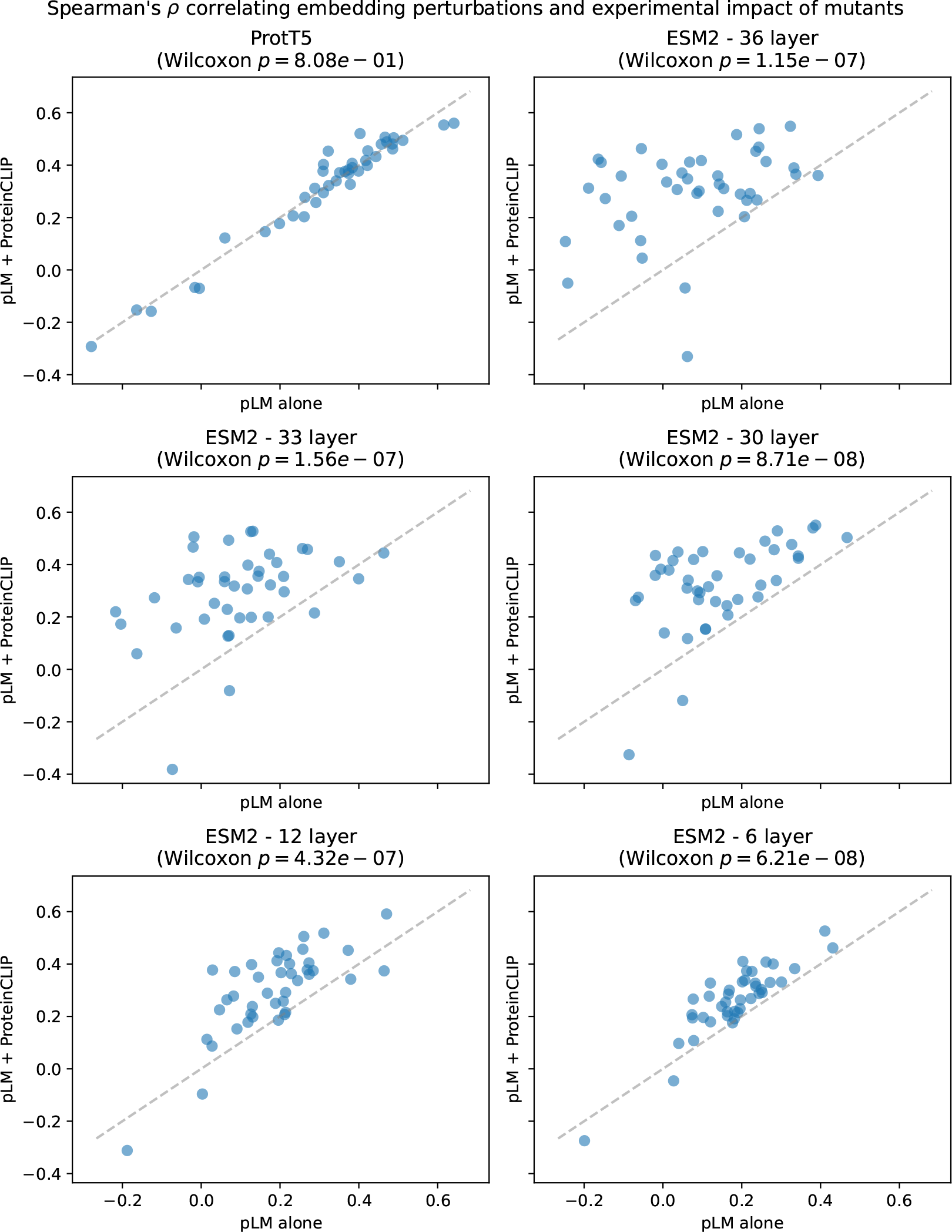
Spearman’s correlation comparing the cosine similarity between a mutated amino acid sequence and wildtype, and the experimentally measured functional impact of that mutation. Each point represents correlation within an experiment. Grey line indicates 1:1 correlation, i.e., equal correlation. The x-axis denotes embedding shift within a pLM’s sequence eembedding, and the y-axis denotes the shift in ProteinCLIP’s sequence embedding. Each panel depicts a different model. All p-values are adjusted for mulitple hypothesis correction using the Holm-Sidak method.

**Supplementary Figure S6:**
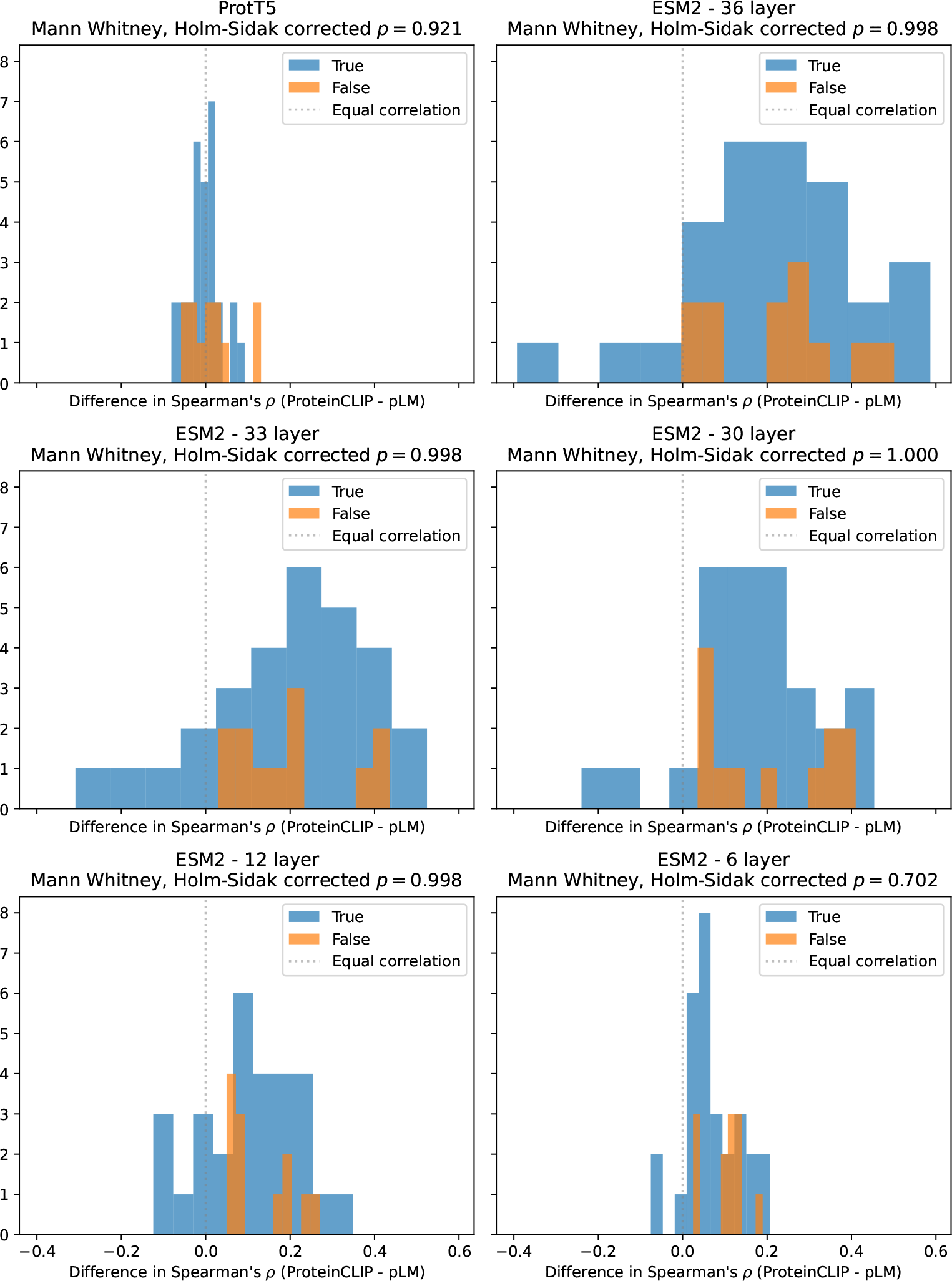
For each of *n* = 41 mutational experiments [27], we calculate the Spearman’s *ρ* between the experimentally measured impact of a mutation and how much that mutation shifts a protein embedding as measured by cosine similarity to reference embedding. Differences between correlations (x-axis) from pLMs and ProteinCLIP models (panels) are plotted as histograms; positive values indicate instances where ProteinCLIP yields higher correlation. These experiments are stratified into those whose wildtype sequence occurs in ProteinCLIP’s training set (blue) and those not seen during training (orange); p-values compare the correlation difference between sequences seen and not seen in training using a Mann-Whitney test and corrected with Holm-Sidak correction. None of the differences are significant.

**Supplementary Figure S7:**
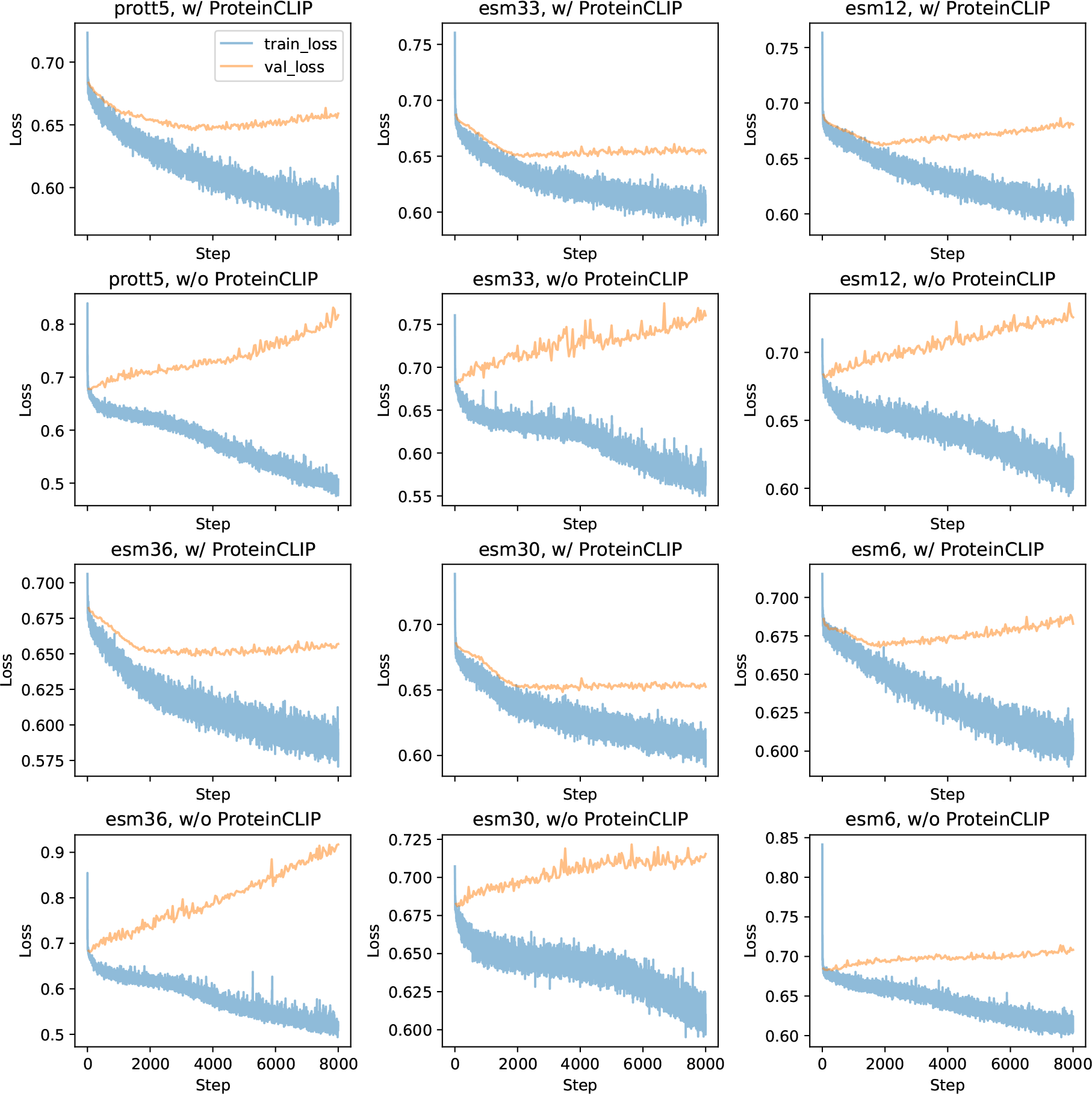
Training curves for protein-protein interaction classifier MLP models. Models are trained for 200 epochs using the “gold standard” training set, allowing sufficient iterations to fit the training set. We compute AUPRC on the “gold standard” validation set to choose the best performing model for further analysis (e.g., evaluation on test data) in each instance.

**Supplementary Figure S8:**
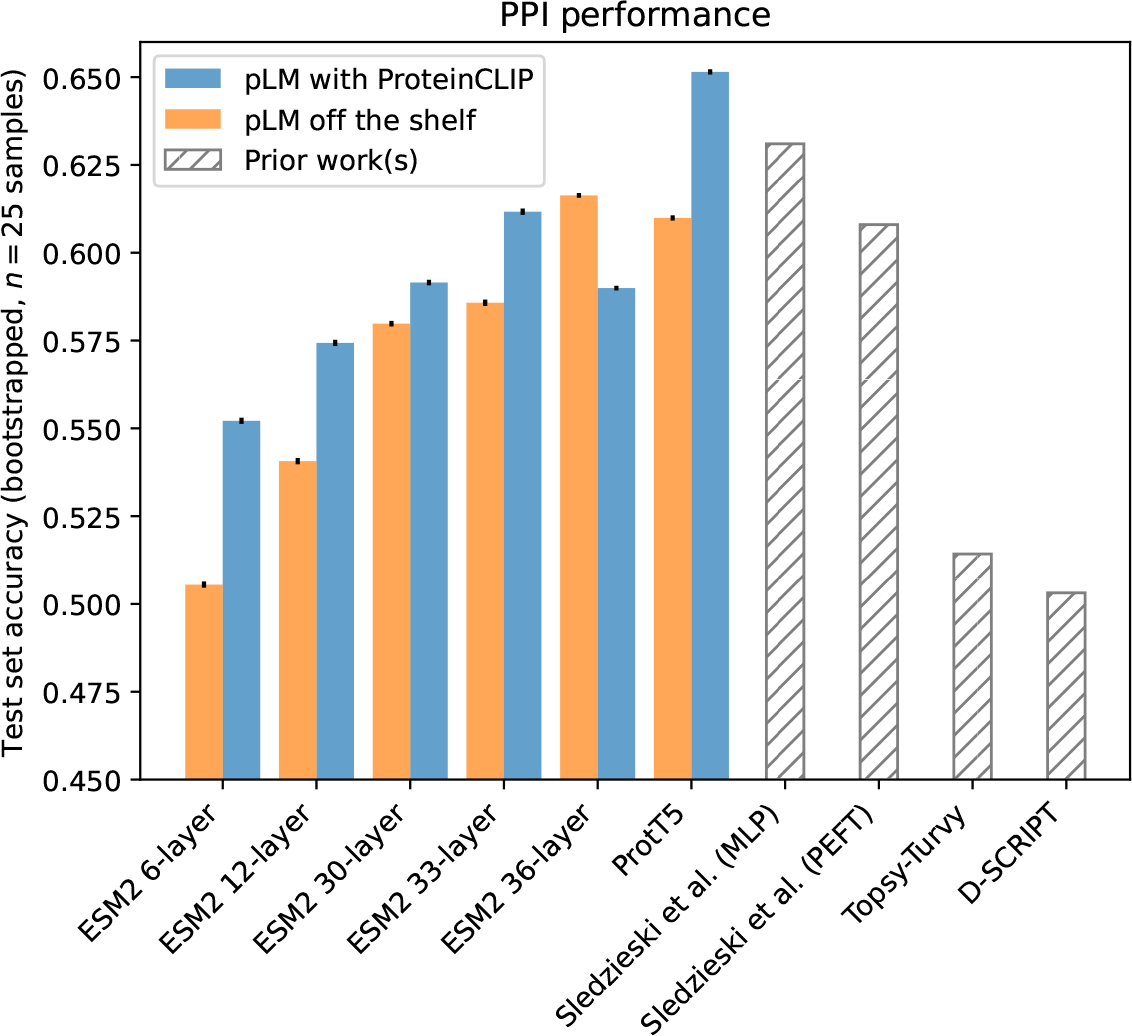
Gold standard test set accuracy of various PPI prediction methods. We evaluate PPI models built directly on pLM embeddings (orange), models built on ProteinCLIP embeddings (blue), and prior works (hatched bars) [35, 34, 19].

**Supplementary Figure S9:**
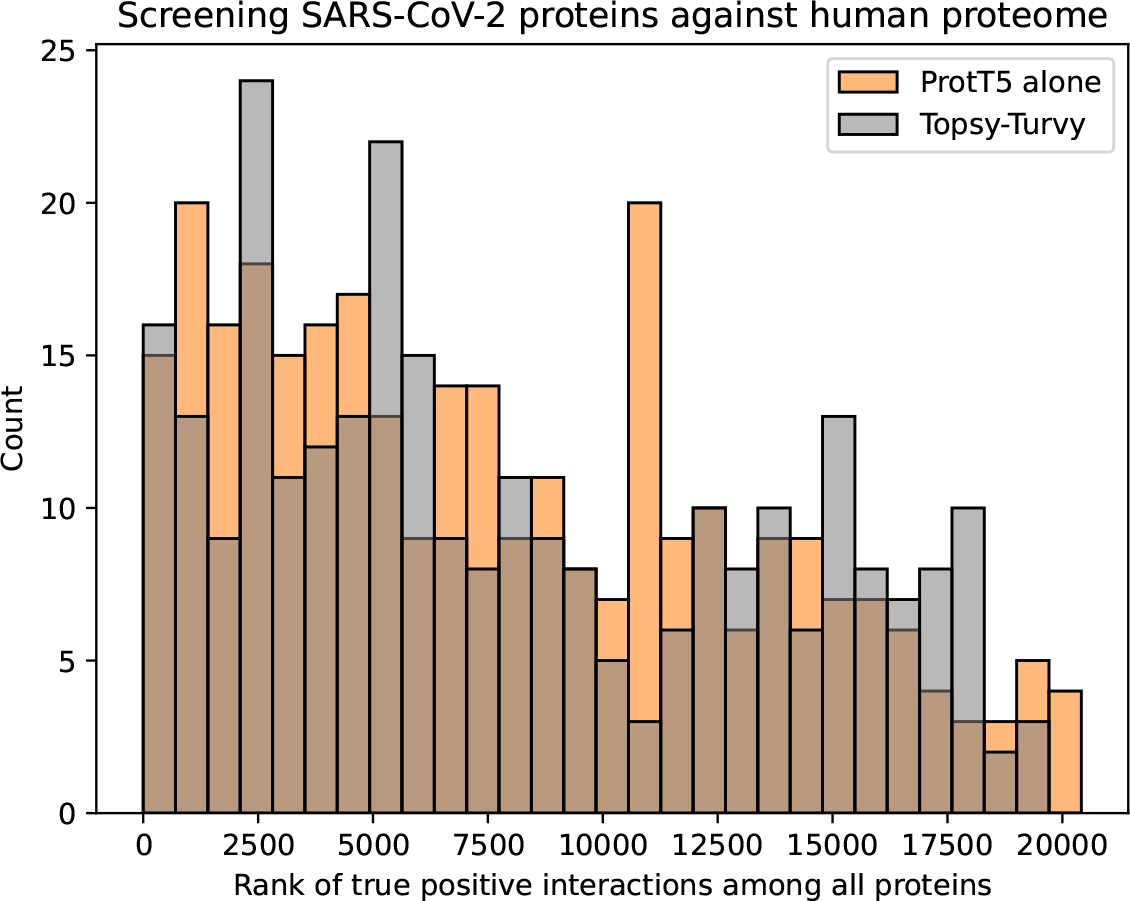
Ranks of true SARS-CoV-2 viral interactions with host human proteins among proteome-wide screen (x-axis) for a classifier built on ProtT5 (without ProteinCLIP, orange) and Topsy-Turvy [34] (grey). There is no significant difference in the distribution of ranks (twosided Wilcoxon test, *p* = 0.320).

**Supplementary Figure S10:**
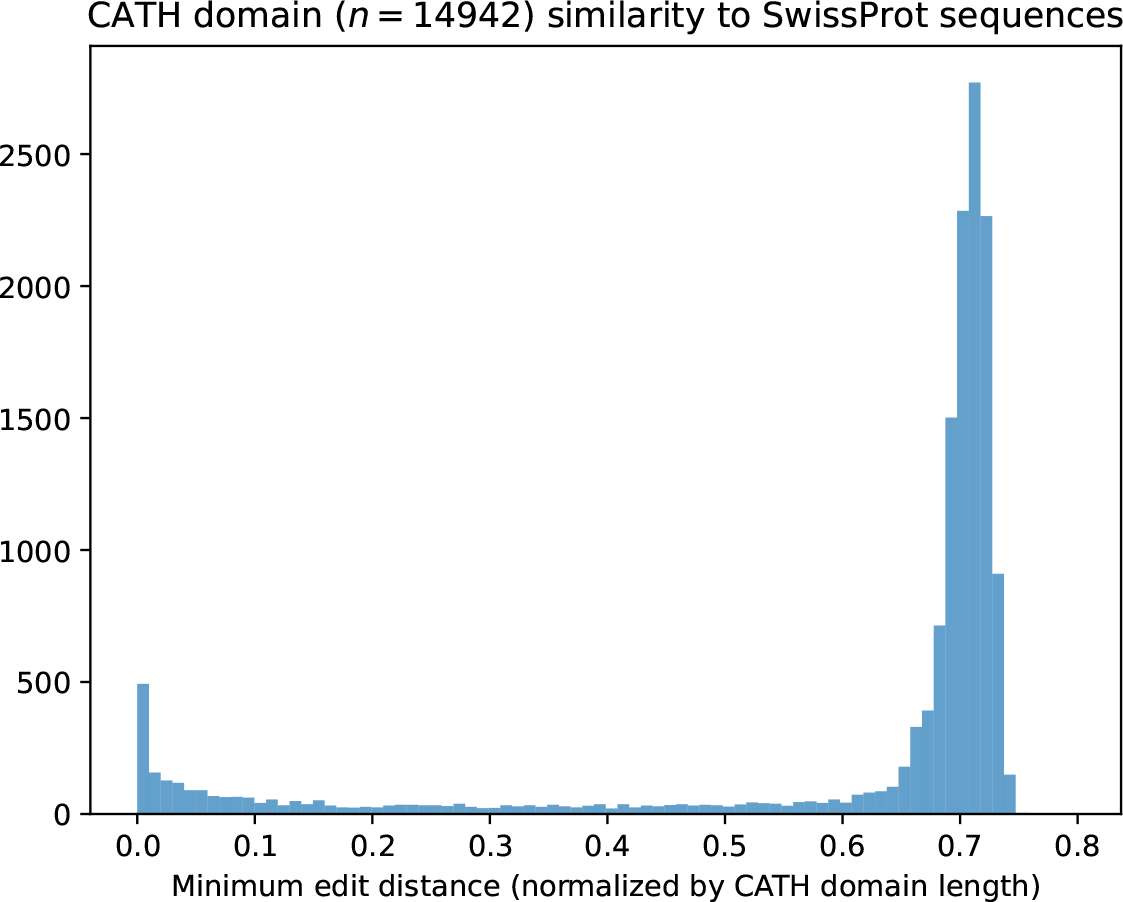
Minimum normalized edit distance of each CATH sequence (*n* = 14942) across all sequences in UniProt, which was used to train ProteinCLIP models. Normalized edit distance is edit distance divided by the length of the CATH sequence in question, and is bound between [0, 1]. 318 (2.1%) of CATH sequences overlap with the UniProt set (i.e., are present exactly), 1335 (8.9%) have a minimum edit distance of 10% or less to any sequence in our UniProt set, and 1715 (11.5%) are have a minimum edit distance of 20% or less.

**Supplementary Table S1:**
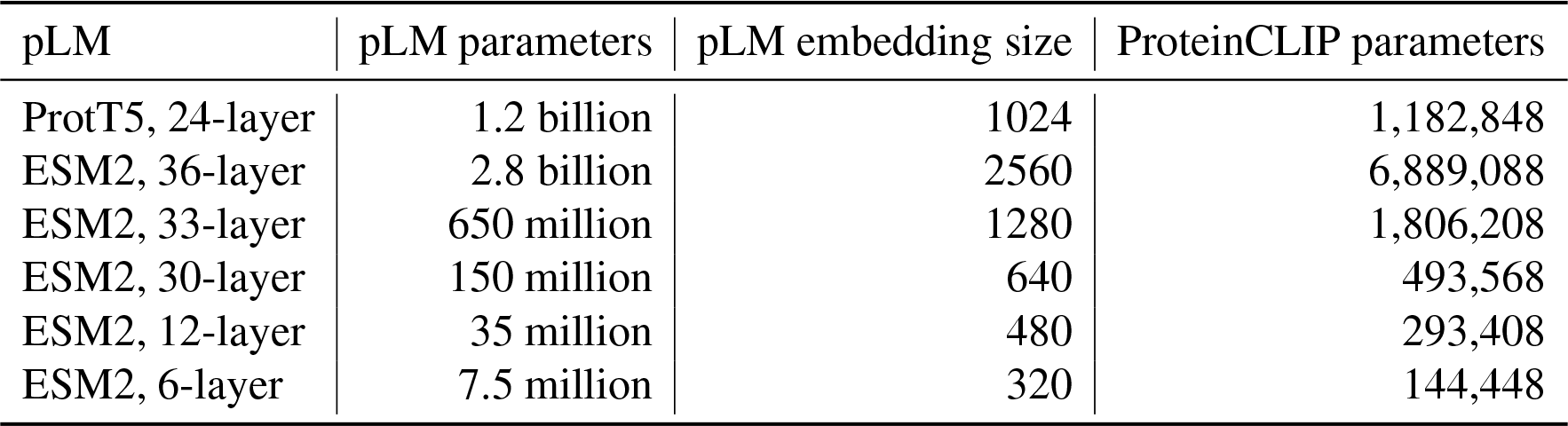
Protein language model (pLM) sizes, embedding dimensions, and corresponding ProteinCLIP model parameter counts. Parameter counts include projection network for the amino acid embeddings, and does not include the projection network for the natural language embedding (as we do not use this network for analyses in this work). All ProteinCLIP models presented project to a final embedding space of *d*_shared_
= 128 dimensions.

| pLM | pLM parameters | pLM embedding size | ProteinCLIP parameters |
| --- | --- | --- | --- |
| ProtT5, 24-layer | 1.2 billion | 1024 | 1,182,848 |
| ESM2, 36-layer | 2.8 billion | 2560 | 6,889,088 |
| ESM2, 33-layer | 650 million | 1280 | 1,806,208 |
| ESM2, 30-layer | 150 million | 640 | 493,568 |
| ESM2, 12-layer | 35 million | 480 | 293,408 |
| ESM2, 6-layer | 7.5 million | 320 | 144,448 |

**Supplementary Table S2:** Key metrics from the function texts used to train ProteinCLIP models. Function texts describe functional attributes of a protein, and do not tend to contain significant metadata attributes like the name of the protein, name of the species the protein is from, or even any interaction partners.

| Category | Number of records |
| --- | --- |
| All records with non-empty function text, $\leq 5800$ residues | 570,037 (100%) |
| Name of protein described (e.g., “LAMP1”) present in function text | 16,895 (3.63%) |
| Name of protein host organism present in function text | 89 (0.0191%) |
| Word “interacts” present in function text (case-insensitive) | 8,148 (1.75%) |

**Supplementary Table S3:** Test set performance on the “gold-standard” PPI prediction dataset proposed by Bernett et al. [33]. Each row shows test set AUPRC, accuracy, and F1-scores for a given PPI classifier model, and the two sub-rows indicate whether or not the classifier’s input embeddings were further augmented by ProteinCLIP (True) or not (False). All PPI classifier models are architecturally the same, with the only difference being the embeddings used as input features. All models were trained on the “gold standard” training dataset with the best model on the “gold standard” validation set measured by AUPRC chosen for further evaluation. For computing accuracy and F1, test set predictions were binarized based on a cutoff determined by computing Youden’s J statistic on the validation set.

| Protein LM | ProteinCLIP | AUPRC | accuracy | F1 |
| --- | --- | --- | --- | --- |
| ProtT5 | True | 0.697 | 0.651 | 0.680 |
|  | False | 0.622 | 0.610 | 0.617 |
| ESM2, 36-layer | True | 0.601 | 0.589 | 0.599 |
|  | False | 0.644 | 0.617 | 0.623 |
| ESM2, 33-layer | True | 0.638 | 0.611 | 0.624 |
|  | False | 0.624 | 0.586 | 0.539 |
| ESM2, 30-layer | True | 0.607 | 0.591 | 0.624 |
|  | False | 0.585 | 0.579 | 0.637 |
| ESM2, 12-layer | True | 0.597 | 0.574 | 0.584 |
|  | False | 0.542 | 0.541 | 0.608 |
| ESM2, 6-layer | True | 0.558 | 0.552 | 0.617 |
|  | False | 0.519 | 0.506 | 0.326 |

**Supplementary Table S4:**
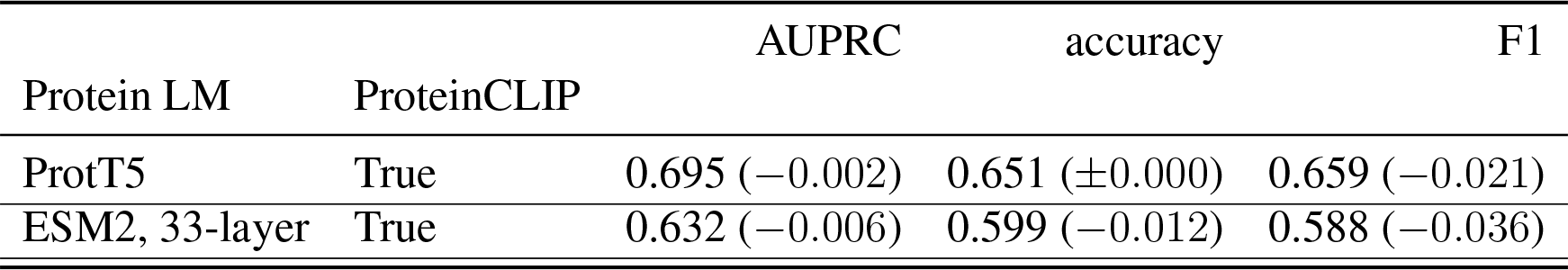
Test set performance on the “gold standard” PPI prediction dataset when both ProteinCLIP and the PPI classifier are trained without seeing any examples in gold standard validation and test sets. Values in parentheses indicate change compared to corresponding model in Table S3.

**Supplementary Table S5:**
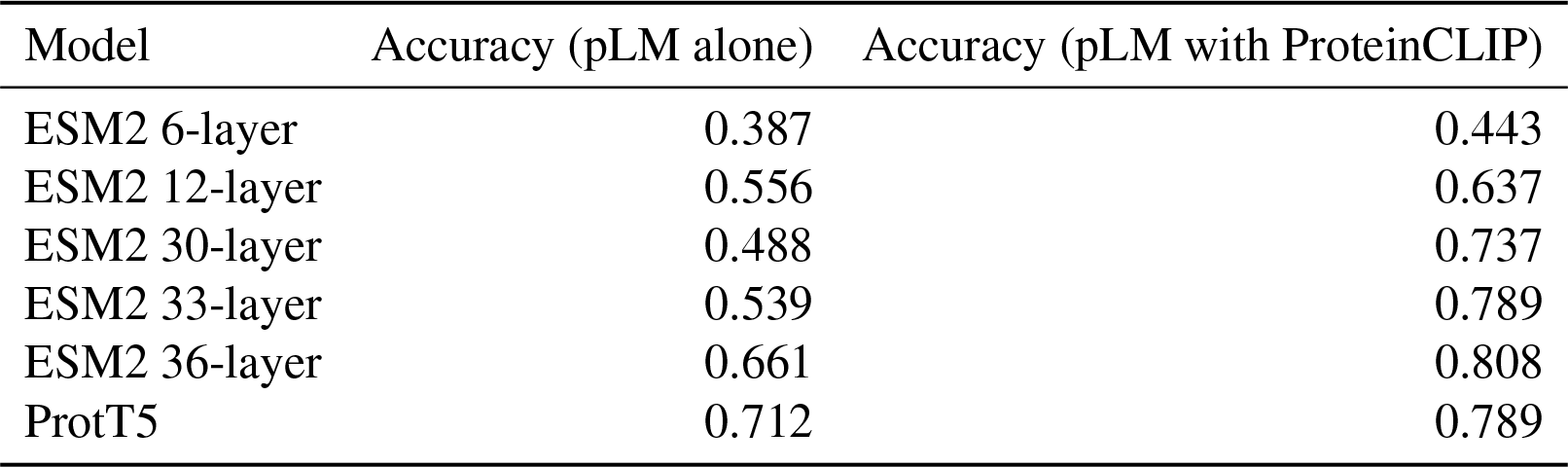
Top-1 accuracy of using different embeddings to retrieve homologous proteins in the CATH S20 dataset. Retrieval is done by fetching the top hit by cosine similarity. Correct hits are defined as sharing the same homologous superfamily as the query sequence. Randomly guessing a superfamily yields an expected accuracy of 1/5559. When evaluated under this same setup, MMseqs2, a state-of-the-art sequence comparison based homology detection method [39], achieves an accuracy of 0.377.

**Supplementary Table S6:**
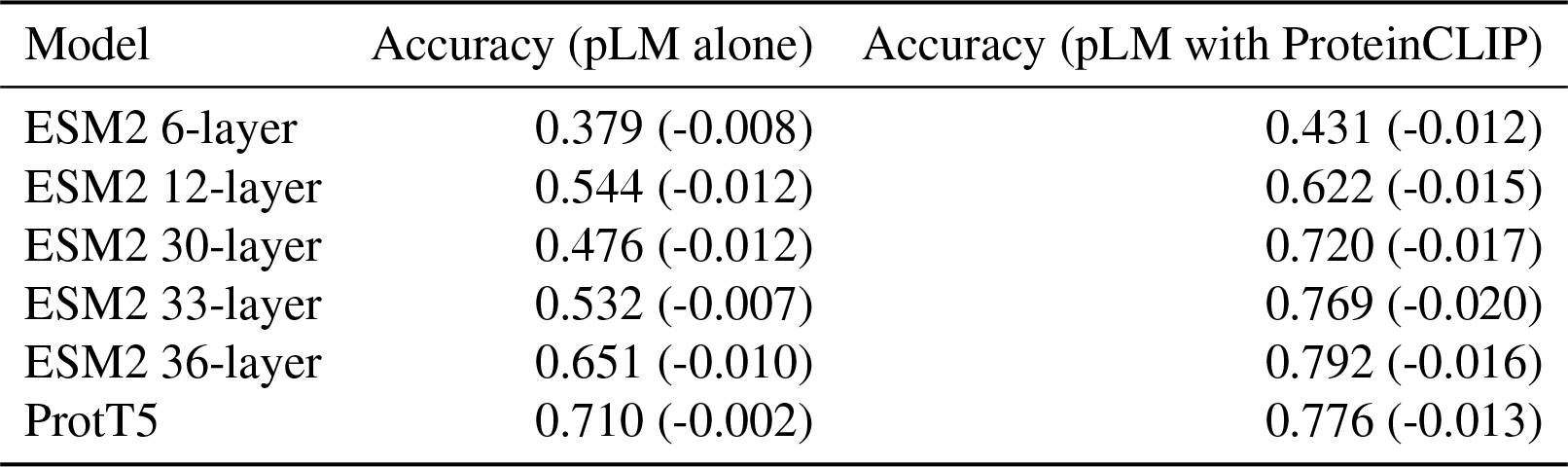
Top-1 accuracy of using different embeddings to retrieve homologous proteins in the CATH S20 dataset, with sequences bearing significant similarity to the UniProt dataset used to train ProteinCLIP removed. Sequences a normalized edit distance of 20% or lower are excluded (Figure S10). Randomly guessing a super family in this setup yields an expected accuracy of 1/5176. Values in parenthesis indicate the change in performance when compared to the same retrieval task performed without sequence similarity removal (Table S5). We never see more than a 2% drop in accuracy under this more stringent evaluation.

## Notes

https://zenodo.org/doi/10.5281/zenodo.11176862

https://github.com/wukevin/proteinclip

### Competing Interest Statement

The authors have declared no competing interest.

